# GPR161 structure uncovers the redundant role of sterol-regulated ciliary cAMP signaling in the Hedgehog pathway

**DOI:** 10.1101/2023.05.23.540554

**Authors:** Nicholas Hoppe, Simone Harrison, Sun-Hee Hwang, Ziwei Chen, Masha Karelina, Ishan Deshpande, Carl-Mikael Suomivuori, Vivek R. Palicharla, Samuel P. Berry, Philipp Tschaikner, Dominik Regele, Douglas F. Covey, Eduard Stefan, Debora S. Marks, Jeremy Reiter, Ron O. Dror, Alex S. Evers, Saikat Mukhopadhyay, Aashish Manglik

## Abstract

The orphan G protein-coupled receptor (GPCR) GPR161 is enriched in primary cilia, where it plays a central role in suppressing Hedgehog signaling^1^. GPR161 mutations lead to developmental defects and cancers^2,3,4^. The fundamental basis of how GPR161 is activated, including potential endogenous activators and pathway-relevant signal transducers, remains unclear. To elucidate GPR161 function, we determined a cryogenic-electron microscopy structure of active GPR161 bound to the heterotrimeric G protein complex G_s_. This structure revealed an extracellular loop 2 that occupies the canonical GPCR orthosteric ligand pocket. Furthermore, we identify a sterol that binds to a conserved extrahelical site adjacent to transmembrane helices 6 and 7 and stabilizes a GPR161 conformation required for G_s_ coupling. Mutations that prevent sterol binding to GPR161 suppress cAMP pathway activation. Surprisingly, these mutants retain the ability to suppress GLI2 transcription factor accumulation in cilia, a key function of ciliary GPR161 in Hedgehog pathway suppression. By contrast, a protein kinase A-binding site in the GPR161 C-terminus is critical in suppressing GLI2 ciliary accumulation. Our work highlights how unique structural features of GPR161 interface with the Hedgehog pathway and sets a foundation to understand the broader role of GPR161 function in other signaling pathways.

## Introduction

Orphan G protein-coupled receptors (GPCRs) coordinate diverse signaling pathways to control many aspects of human physiology^5^. The orphan GPCR GPR161 has been characterized as a unique example of a constitutively active receptor that is located within the primary cilium of cells, an organelle that protrudes from the cell surface and locally organizes signaling components^1^. In its best understood signaling role, GPR161 is a critical negative regulator of the Hedgehog pathway^1^. Knockout of *Gpr161* in mice is embryonically lethal, and the embryos display severe limb, facial, and early nervous system defects indicative of hyperactive Hedgehog signaling^6–10^. *GPR161* mutations in humans lead to developmental defects such as spina bifida^2,11,12^, pituitary stalk interruption syndrome^13^, and cancers such as medulloblastoma^3,7^. Overexpression of GPR161 has been linked to triple-negative breast cancer^4^. Like many other orphan GPCRs, however, fundamental mechanisms of GPR161 function remain unknown^14^, including what stimulus gives rise to GPR161 constitutive activity and how signaling activity downstream of GPR161 impinges on its biological function.

The primary function of GPR161 has been framed by its discovery as a Hedgehog pathway regulator^1^. Hedgehog signaling during vertebrate embryogenesis mediates multicellular development, including the proper formation of limbs, the face, and the nervous system^15^. In the presence of the Hedgehog signal, GLI2/3 transcriptional factors accumulate in the primary cilia and form activators (GLI-A)^16^. In the absence of the Hedgehog signal, GLI2/3 are constitutively phosphorylated by protein kinase A (PKA), which leads to proteolytic conversion of these proteins into Hedgehog pathway repressors (GLI-R). Because PKA is canonically activated by the GPCR second messenger cyclic adenosine monophosphate (cAMP), current models propose that elevated ciliary cAMP levels activate PKA to suppress the Hedgehog pathway^17^. Although many GPCRs localize to the primary cilium^18,19^, several observations have placed GPR161 as a unique Hedgehog pathway regulator. Loss of GPR161 function in mice and in fish causes phenotypes consistent with inappropriate Hedgehog pathway activation^6–10,20^. GPR161 functions both inside cilia and in the peri-ciliary endosomal compartments in regulating these phenotypes^1,6^. Furthermore, GPR161 is constitutively active in model cell lines and drives elevated cAMP via activation of G_s_^1,6,20–22,23,24^. Upon Hedgehog pathway activation, GPR161 exits cilia by internalizing to the recycling endocytic compartment^1,21^. Finally, the C-terminus of GPR161 binds specifically PKA type I regulatory subunits through an A-kinase anchoring protein domain (AKAP), a unique feature of GPR161 among hundreds of GPCRs^25^. GPR161 is therefore thought to repress Hedgehog signaling by constitutive coupling to G_s_, which elevates cAMP levels to drive PKA activity (Fig. 1a).

**Figure 1:**
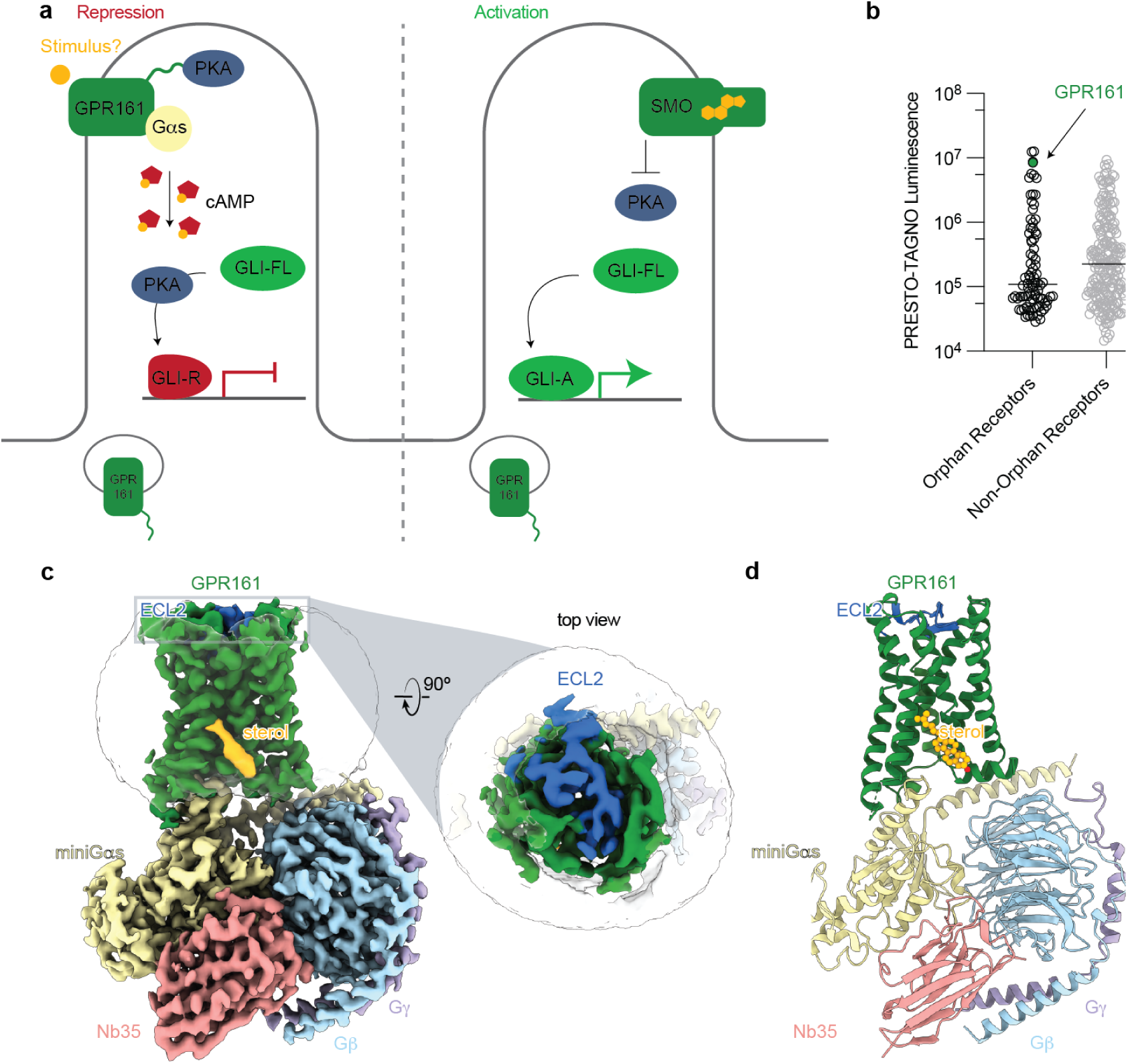
Structure-inspired deorphanization of GPR161. **a)** The Hedgehog pathway is regulated by two key GPCRs, GPR161 and SMO. In the absence of Hedgehog, GPR161 represses the pathway through constitutive cAMP signaling from an unknown stimulus. In the presence of Hedgehog, SMO activates the pathway by entering the cilia, binding cholesterol, and inhibiting PKA, while GPR161 exits cilia. **b)** GPR161 yields an exceptionally strong signal for β-arrestin recruitment in the PRESTO-Tango assay when compared across 314 GPCRs (data replotted from Kroeze WM *et al* 2015^22^). This assay is performed in a modified HEK293 cell line, suggesting that GPR161 is constitutively active under heterologous expression conditions. **c)** Cryo-EM density map of GPR161 in complex with G_s_ heterotrimer (miniGα_s_, Gβ, and Gγ) and stabilizing nanobody 35 (Nb35). The map reveals a density consistent with the shape of a sterol (yellow) and an extracellular loop 2 (ECL2, blue) that is packed within the seven transmembrane core of GPR161. **d)** Ribbon diagram of activated GPR161 heterotrimer complex. Cholesterol is modeled into the sterol density (yellow).

Several fundamental aspects of GPR161 function remain unclear - in particular, the potential stimulus that drives GPR161 activity remains unknown. The interdependent roles of G_s_ coupling and PKA binding, and their relative importance in Hedgehog signaling is also poorly defined. Here we use a combination of cryogenic-electron microscopy (cryo-EM), signaling studies, molecular dynamics simulations, and biochemical assays to determine the molecular mechanism of GPR161 activation and Hedgehog pathway repression. Our studies reveal that GPR161-induced G_s_ signaling is driven by a novel sterol-binding site. However, this signaling activity does not repress GLI2 ciliary accumulation, a key role of ciliary GPR161 in Hedgehog pathway repression. By contrast, the AKAP domain in GPR161 is necessary for repressing GLI2 accumulation in cilia. Together, these findings provide an activation mechanism for GPR161 and support PKA as a central downstream ciliary regulator of the Hedgehog pathway.

### Cryo-EM structure of active GPR161 bound to G_s_ heterotrimer

GPR161 is one of the most constitutively active GPCRs tested in the β-arrestin PRESTO-Tango assay^22^, which uses nonciliated HEK293 cells (Fig. 1b). We reasoned that purification of GPR161 from HEK293 cells may allow us to determine a structure of the active signaling state and may reveal potential stimuli for GPR161. Preparations of GPR161 in the absence of a signal transducer were of poor quality, suggesting that GPR161 alone may be structurally dynamic or otherwise unstable. To stabilize GPR161 in the active state and simultaneously increase the likelihood that the receptor would co-purify with an activating stimulus^26^, we C-terminally fused the receptor to a minimal version of the Gα_s_ protein. This minimized “miniG_s_” construct retains the receptor-interacting GTPase domain of the Gα_s_ subunit but is engineered to interact with a GPCR in the absence of a guanine nucleotide^27^. We purified GPR161-miniG_s_ to homogeneity, further complexed it with other heterotrimeric G protein subunits Gβ_1_ and Gγ_2_ as well as nanobody 35 (Nb35) to stabilize the interaction between Gα_s_ and Gβ_1_γ_2_ (Supplementary Fig. 1)^28^. The resulting complex was imaged by cryogenic-electron microscopy to yield a reconstruction of the GPR161-G_s_ complex at 2.7 Å resolution (Fig. 1c,d and Supplementary Fig. 2) and enabled model building for the seven transmembrane domain of GPR161, the G_s_ subunits, Nb35, and, most notably, a sterol-like molecule (Fig. 1c,d and Supplementary Fig. 3).

Our structure of the GPR161-G_s_ complex is similar to many other activated Class A GPCRs bound to heterotrimeric G proteins, like the prototypical β_2_-adrenergic receptor (β_2_AR) (Supplementary Fig. 4a). A key hallmark of Class A GPCR activation is outward displacement of transmembrane helix 6 (TM6) to accommodate the C-terminal α-helix of Gα subunit^29^. While we do not have an inactive structure of GPR161 for comparison, the conformation of TM6 in GPR161 bound to G_s_ is similar to the outward displaced conformation observed for β_2_AR (Supplementary Fig. 4a). We conclude that our structure of GPR161-G_s_ captures the G_s_ coupled, active conformation of the receptor.

### GPR161 extracellular loop 2 is self-activating

The structure of active GPR161 revealed a unique conformation of extracellular loop 2 (ECL2) compared to the majority of ligand-activated Class A GPCRs. Notably, the ECL2 of GPR161 forms a beta hairpin that folds over the extracellular face of the receptor to completely occlude the canonical orthosteric ligand binding pocket observed for many other Class A GPCRs (Fig. 1c,d and Fig. 2a). The comparison to β_2_AR, for example, highlights that the GPR161 ECL2 occupies the same space that adrenaline occupies in β_2_AR (Fig. 2b). The conformation of ECL2 in GPR161 is reminiscent of the orphan GPCR GPR52, which contains an ECL2 that also occludes the extracellular face of the receptor (Fig. 2b)^30^. In GPR52, ECL2 serves as a key determinant of constitutive activity - in effect, GPR52 is “self-activated” by ECL2. Indeed structures of several other orphan GPCRs, including GPR21 and GPR17 have recently revealed similar conformations of ECL2 associated with self-activation (Supplementary Fig. 4b)^31,32^.

**Figure 2:**
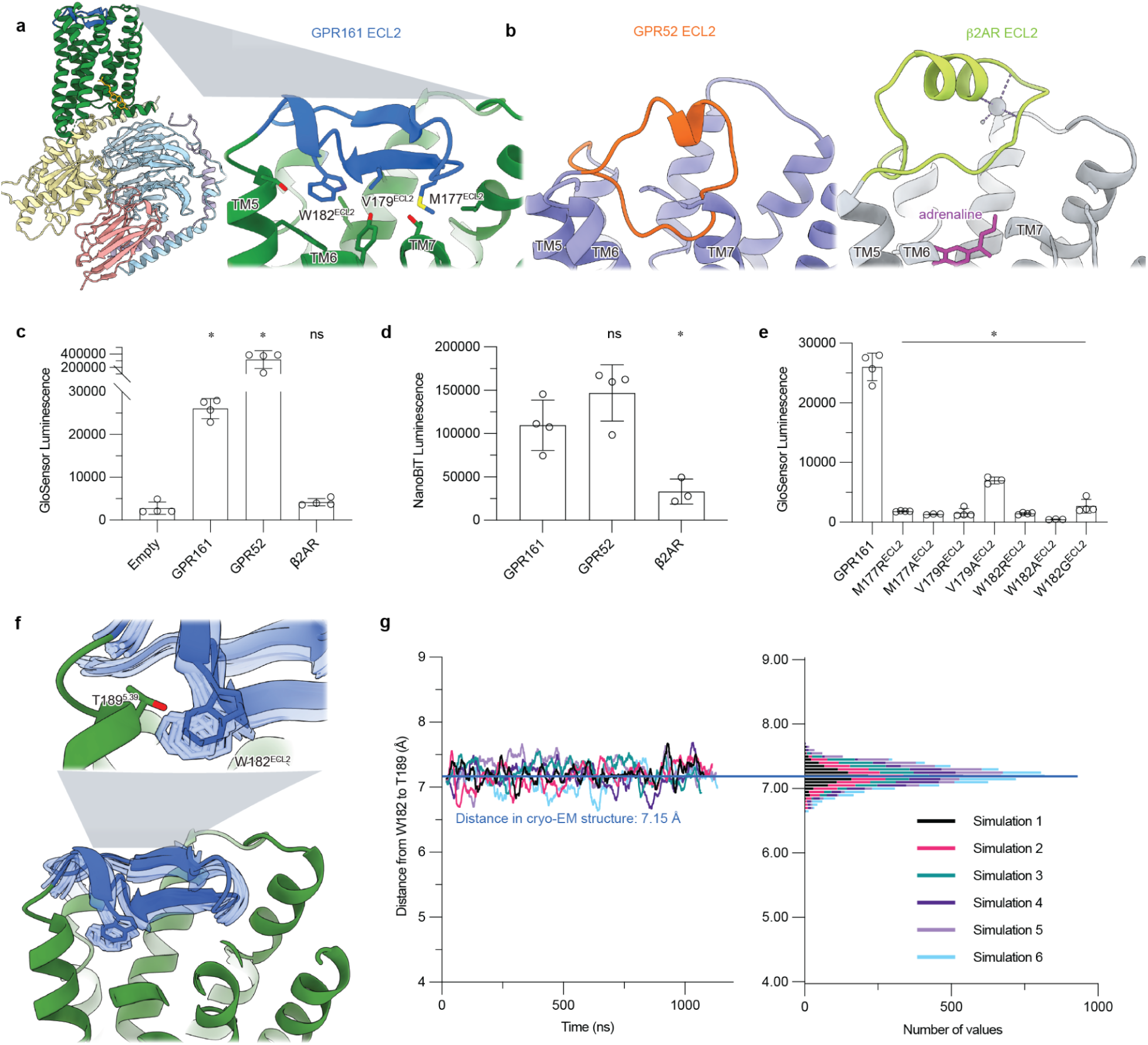
Extracellular loop 2 of GPR161 occupies classic GPCR orthosteric site. **a**) The ECL2 of GPR161 makes hydrophobic contacts with the core of the receptor. **b)** Comparison of ECL2 of the self-activating orphan GPCR GPR52 (PDB ID: 6LI3^30^) and the prototypical agonist-activated GPCR β2-adrenergic receptor (β_2_AR) bound to agonist adrenaline (PDB ID: 4LDO^65^). **c)** cAMP production assay for GPR161, GPR52, and β_2_AR. GPR161 is constitutively active for cAMP production. Data are mean ± sd, n=4 biologically independent samples (\**P* < 0.05; ns, not significant; one-way ANOVA followed by Dunnett’s multiple comparison tests). **d)** Nanoluc complementation assay for receptor recruitment of miniG_s_. Both GPR52 and GPR161 constitutively recruit miniG_s_. Data are mean ± sd, n=3-4 biologically independent samples (\**P* < 0.05; ns, not significant; one-way ANOVA followed by Dunnett’s multiple comparison tests). **e)** cAMP production assay assessing mutations in ECL2 of GPR161 for residues that make hydrophobic contacts with the transmembrane bundle. Data are mean ± sd, n=4 biologically independent samples (\**P* < 0.05; ns, not significant; one-way ANOVA followed by Dunnett’s multiple comparison tests). **f)** Molecular dynamics simulations snapshots of GPR161. **g)** ECL2 stably occupies the canonical ligand binding pocket as represented by rolling average of the distance between ECL2 W182^ECL2^ and the Cα atom of T189^5.39^ in molecular dynamics simulations (6 independent trajectories).

To first verify the constitutive activity of GPR161, we used two different cellular assays: GloSensor assay to assess cAMP production and a miniG_s_ recruitment assay^33^ using an optimized NanoLuciferase fragment complementation termed “NanoBiT”^34^. Expression of GPR161 in suspension adapted HEK293 cells gave consistently high levels of cAMP under basal conditions relative to empty vector and β_2_AR (13 and 8 fold, respectively) (Fig. 2c and Supplementary Fig. 5). However, GPR161 produced markedly less cAMP than GPR52 (Fig. 2c), perhaps highlighting that self-interaction of ECL2 is not sufficient to drive high basal activity. In a miniG_s_ protein fragment complementation assay, GPR161 basally recruited more miniG_s_ than β_2_AR, with levels that are more similar to GPR52 (Fig. 2d). The results from these two orthogonal assays demonstrate that GPR161 is constitutively active, albeit to a lesser extent than the self-activating orphan receptor GPR52.

Inspired by the example of GPR52, we examined the possible role of ECL2 in GPR161 activation. Several hydrophobic residues of the GPR161 ECL2 protrude into a region that overlaps with the canonical orthosteric pocket in other Class A GPCRs (Fig. 2a). We targeted several of these (M177^ECL2^, V179^ECL2^, W182^ECL2^) for mutagenesis experiments to understand whether the conformation of ECL2 in GPR161 is important for constitutive activity. We substituted each of these positions with either alanine (to test for simple loss of the side chain) or arginine (to introduce a large perturbation in local hydrophobic contacts). We also examined a W182G^ECL2^ mutant, which has previously been associated with rare cases of spina bifida, a neural tube developmental defect^35^. Mutation of these hydrophobic residues in ECL2 to either alanine or arginine caused a near complete loss of cAMP generation by GPR161, suggesting that the in *cis* interaction with ECL2 is essential for GPR161 activation (Fig. 2e). To explore the possibility that the GPR161 ECL2 may be more dynamic in the absence of miniG_s_, we performed molecular dynamics simulations of GPR161 without miniG_s_. In six simulations, we observed that ECL2 remains stably in a similar conformation as observed in the cryo-EM structure (Fig. 2f,g and Supplementary Fig. 8b). We therefore surmise that ECL2 contributes to self-activation of GPR161.

### A sterol facilitates GPR161 coupling to G_s_

A surprising finding in the cryo-EM map of GPR161-G_s_ is the presence of a sterol-like density located at an extrahelical site near the cytoplasmic ends of transmembrane helices 6 and 7 (TM6 and TM7). Although the exact identity of this sterol is unclear, we tentatively modeled a cholesterol molecule in this density. We next sought to understand whether sterols engage this site and whether sterol binding at this site leads to signaling output for GPR161. Given the importance of sterols in metazoan Hedgehog pathway signaling^36–38^, we first examined whether residues surrounding the putative sterol are conserved in evolution. Several key interacting residues (I323^7.52^, W327^7.56^ and R332^8.51^) are conserved from humans to the echinoderm *Strongylocentrotus purpuratus* (Fig. 3b and Supplementary Fig. 6b).

**Figure 3:**
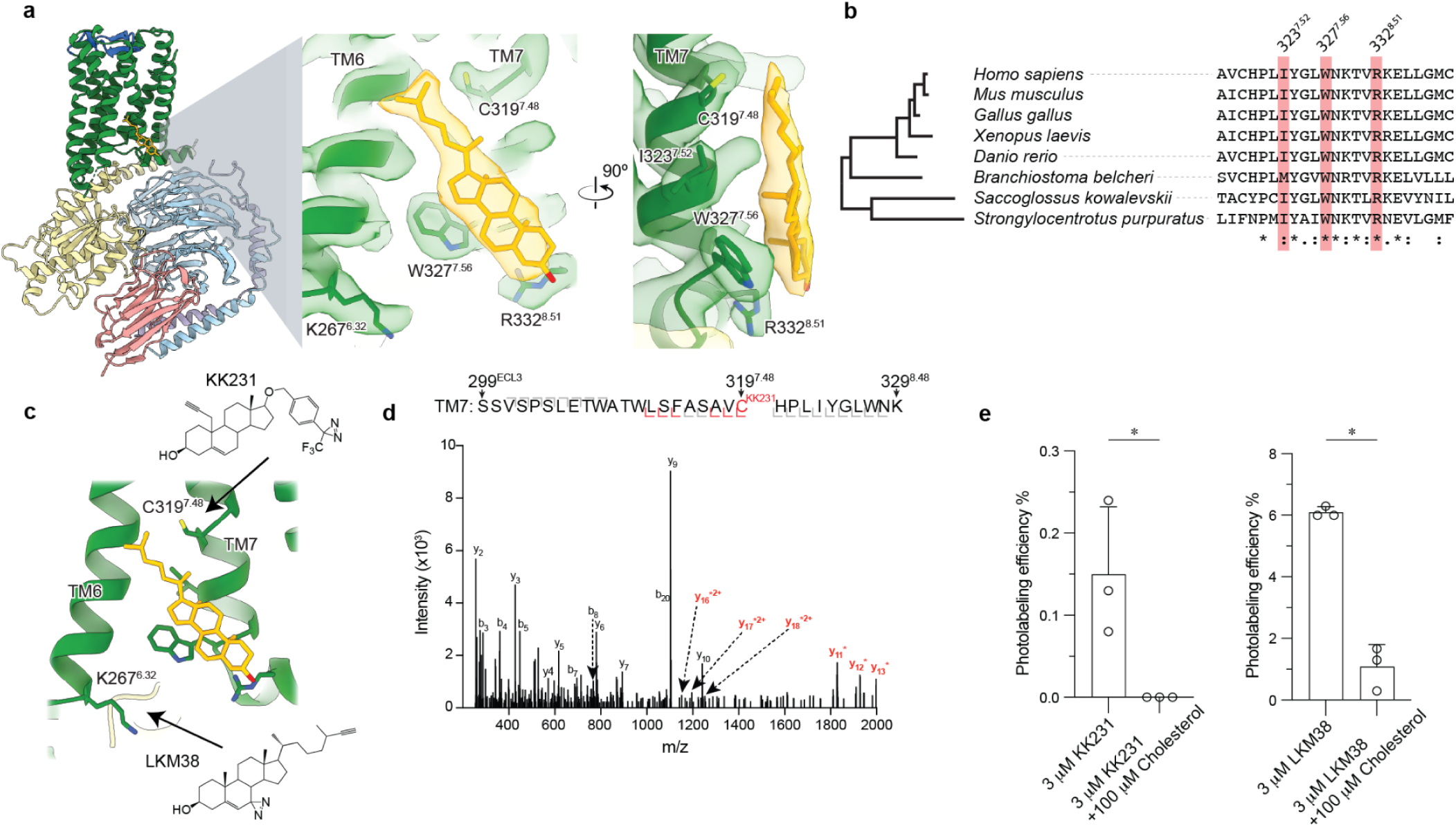
GPR161-miniG_s_ stably and specifically binds cholesterol. **a)** Close up view of cholesterol bound to the intracellular side of transmembrane helix 6 (TM6) and TM7. Three key interacting residues (I323^7.52^, W327^7.56^, R332^8.51^) are highlighted as sticks. **b)** GPR161 cholesterol binding site residues are conserved from humans to sea urchins (*Strongylocentrotus purpuratus*). A full alignment from these organisms used to define the dendrogram on left is shown in Supplementary Fig. 6. **c)** Detergent solubilized GPR161-miniG_s_ purified without cholesterol hemisuccinate was incubated with either one of two distinct photoaffinity cholesterol analogs (KK231 or LKM38) and then crosslinked with >320 nm UV light. The resulting photo-labeled preparation was digested with trypsin and analyzed by collision-induced dissociation mass spectrometry, which revealed that KK231 labels position C319^7.48^ while LKM38 labels K267^6.32^. **d)** Product ion spectrum of KK231-labeled GPR161-miniG_s_ sample with peptides mapped to TM7 and Helix 8. This peptide is modified with a mass consistent with KK231 at position C319^7.48^. Red brackets and peaks indicate product ions that contain the KK231 adduct. **e)** Photolabeling efficiency of GPR161-miniG_s_ by KK231 and LKM238 in the absence or presence of excess unlabeled cholesterol. Data are mean ± s.d of *n*=3 technically independent replicates from two independently prepared protein samples (\**P* < 0.05, Student’s paired t-test).

To determine whether the observed density is indeed a sterol, we performed photoaffinity labeling experiments with purified GPR161-miniG_s_ and two cholesterol analogs: LKM38 and KK231. These sterol analogs contain an ultraviolet light activated diazirine group, either on the B-ring of the steroid (LKM38) or the aliphatic tail (KK231), that rapidly forms covalent adducts with proximal residues. Previous studies have demonstrated that these sterol photoaffinity analogs enable identification of functionally-relevant sterol binding sites in diverse membrane proteins^39–42^. After photoaffinity labeling GPR161-miniG_s_, adducted residues were identified by tryptic digestion followed by LC-MS/MS sequencing of the resultant peptides. We obtained 76% sequence coverage of GPR161, with full residue-level sequencing of six of the seven transmembrane helices (Supplementary Fig. 7b). Consistent with our binding pose for cholesterol, the diazirine group in the B ring of LKM38 labeled K267^6.32^ in TM6 while the similar functional group in the tail of KK231 labeled residue C319^7.48^ in TM7 (Fig. 3c,d and Supplementary Fig. 7a). To determine whether the observed photolabeling is specific, we repeated this experiment in the presence of a 33-fold molar excess of unlabeled cholesterol. For both LKM38 and KK231 the presence of unlabeled cholesterol completely suppressed photoaffinity labeling, suggesting that cholesterol itself can bind at this site (Fig. 3e).

We next sought to understand whether cholesterol binding promotes interactions between GPR161 and G_s_. In the absence of an inactive-state structure of GPR161, we turned to molecular dynamics simulations to assess whether the presence of G_s_ is required for stable cholesterol binding (Fig. 4a,b and Supplementary Fig. 8a). We simulated GPR161 either restrained to remain in the miniG_s_ conformation on the intracellular side or without any restraints. Each condition was simulated with six replicate simulations, each 1 µs in length. When GPR161 is restrained in the miniG_s_ bound conformation, 5 of 6 simulations showed stable cholesterol association with W327^7.56^ (Supplementary Fig. 8d). By contrast, in the absence of any restraints, W327^7.56^ flipped inward into the seven transmembrane core of GPR161, thereby removing a key contact for cholesterol at the extrahelical binding site. Additionally, this rotamer of W327^7.56^ would occlude binding of the C-terminal α-helix of G_s_ (Supplementary Fig. 8e). Indeed, in 5 of 6 simulations of unrestrained GPR161, we observed that cholesterol rapidly disengages the extrahelical binding site and remains unbound for the remainder of the simulation (Supplementary Fig. 8c). GPR161 did not transition to an inactive-like conformation in unrestrained simulations. These simulations therefore suggested that cholesterol binding to GPR161 at the TM6/TM7 extrahelical site is cooperative with G_s_ binding.

**Figure 4:**
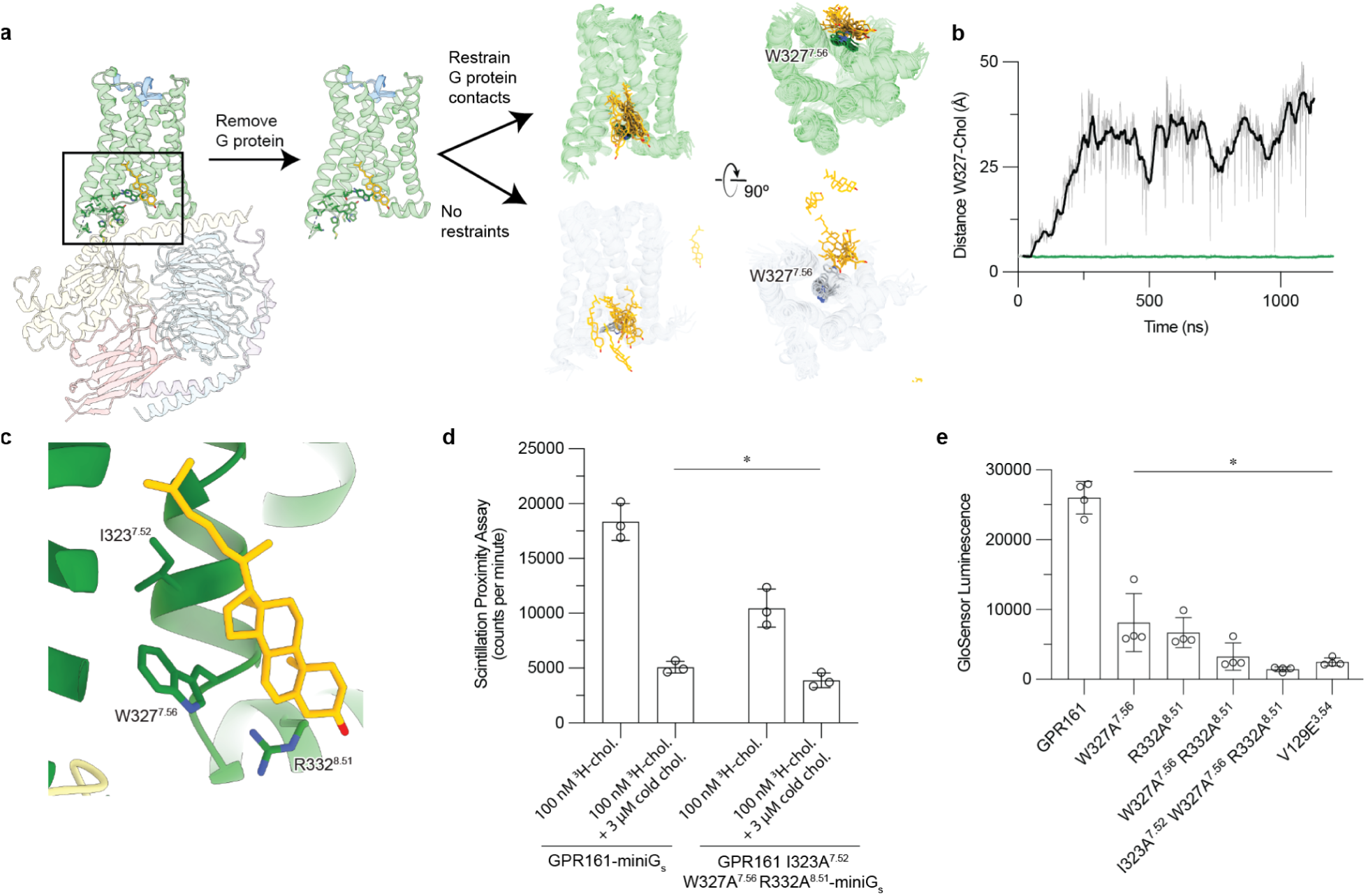
Cholesterol binding to GPR161 facilitates G_s_ coupling. **a)** Molecular dynamics simulation snapshots of active GPR161 both unrestrained (gray) and restrained to G protein bound conformation (green). When GPR161 is restrained to be in the G protein-bound conformation, cholesterol remains consistently near W327^7.56^ as shown in both simulation snapshots and in a time trajectory from a representative simulation. **b)** By contrast, when GPR161 is not restrained to be in the G protein-bound state, the cholesterol is dynamic and moves away from W327^7.56^. **c)** Close up view of cholesterol bound to the intracellular side of transmembrane helix 6 (TM6) and TM7. Three key interacting residues (I323^7.52^, W327^7.56^, R332^8.51^) are highlighted as sticks. **d)** ^3^H-cholesterol (^3^H-chol) binding assay for purified GPR161-miniG_s_ and GPR161-AAA^7.52, 7.56, 8.51^-miniG_s_. Individual technical replicates are shown, bar graphs and error represent mean and s.d. (\**P* < 0.05; two-way ANOVA followed by Dunnett’s multiple comparison tests). **e)** GloSensor cAMP production assay assessing mutations in cholesterol binding site of GPR161. Data are mean ± s.d. from *n*=4 biologically independent replicates (\**P* < 0.05; one-way ANOVA followed by Dunnett’s multiple comparison tests).

If cholesterol potentiates G_s_ binding, we predicted that disrupting the cholesterol binding site would both decrease cholesterol binding and may decrease cAMP production downstream of GPR161-induced G_s_ activation. Closer examination of the conserved residues in the sterol site highlighted that I323^7.52^ interacts with the iso-octyl tail of cholesterol, W327^7.56^ binds cholesterol central rings, and R332^8.51^ engages the hydroxyl group (Fig. 4c). We generated a GPR161-miniG_s_ construct substituting alanine at these conserved positions (GPR161-AAA^7.52,7.56, 8.51^-miniG_s_) and tested the ability of purified receptor preparations to bind ^3^H-cholesterol using a scintillation proximity assay (Fig. 4d). In this assay, GPR161-AAA^7.52, 7.56, 8.51^-miniG_s_ bound cholesterol less than as compared with GPR161-miniG_s_. We observe some residual binding of ^3^H-cholesterol to GPR161-AAA^7.52, 7.56, 8.51^-miniG_s_, suggesting that I323^7.52^, W327^7.56^ and R332^8.51^ help mediate the sterol interaction. Residual binding of ^3^H-cholesterol may reflect GPR161-AAA^7.52, 7.56, 8.51^-miniG_s_ being trapped by miniG_s_ in the G_s_-interacting conformation. Our attempts to conduct this binding experiment with GPR161 alone was limited by the inability to purify the receptor in the absence of miniG_s_. Supporting the importance of the sterol binding site, alanine mutation of W327^7.56^ and R332^8.51^ showed decreased cAMP production in a GloSensor assay compared to wild-type, while the GPR161-AAA^7.52, 7.56, 8.51^ mutant ablates cAMP production (Fig. 4e). Importantly the GPR161-AAA^7.52, 7.56, 8.51^ mutant showed even lower levels of cAMP production compared to V129E^3.54^, a mutant previously designed to directly disrupt the predicted GPR161-G_s_ interaction^1^. Similarly, GPR161-AAA^7.52, 7.56, 8.51^ mutant did not recruit miniG_s_ in a NanoLuciferase complementation assay^33^ (Fig. 5d) while GPR161-V129E showed a more moderate decrease in miniG_s_ recruitment (Supplementary Fig. 9d). Our combined biochemical, simulation, and signaling studies show that cholesterol, and potentially other sterols, can bind GPR161 to support interactions with G_s_, thereby promoting cAMP production.

**Figure 5:**
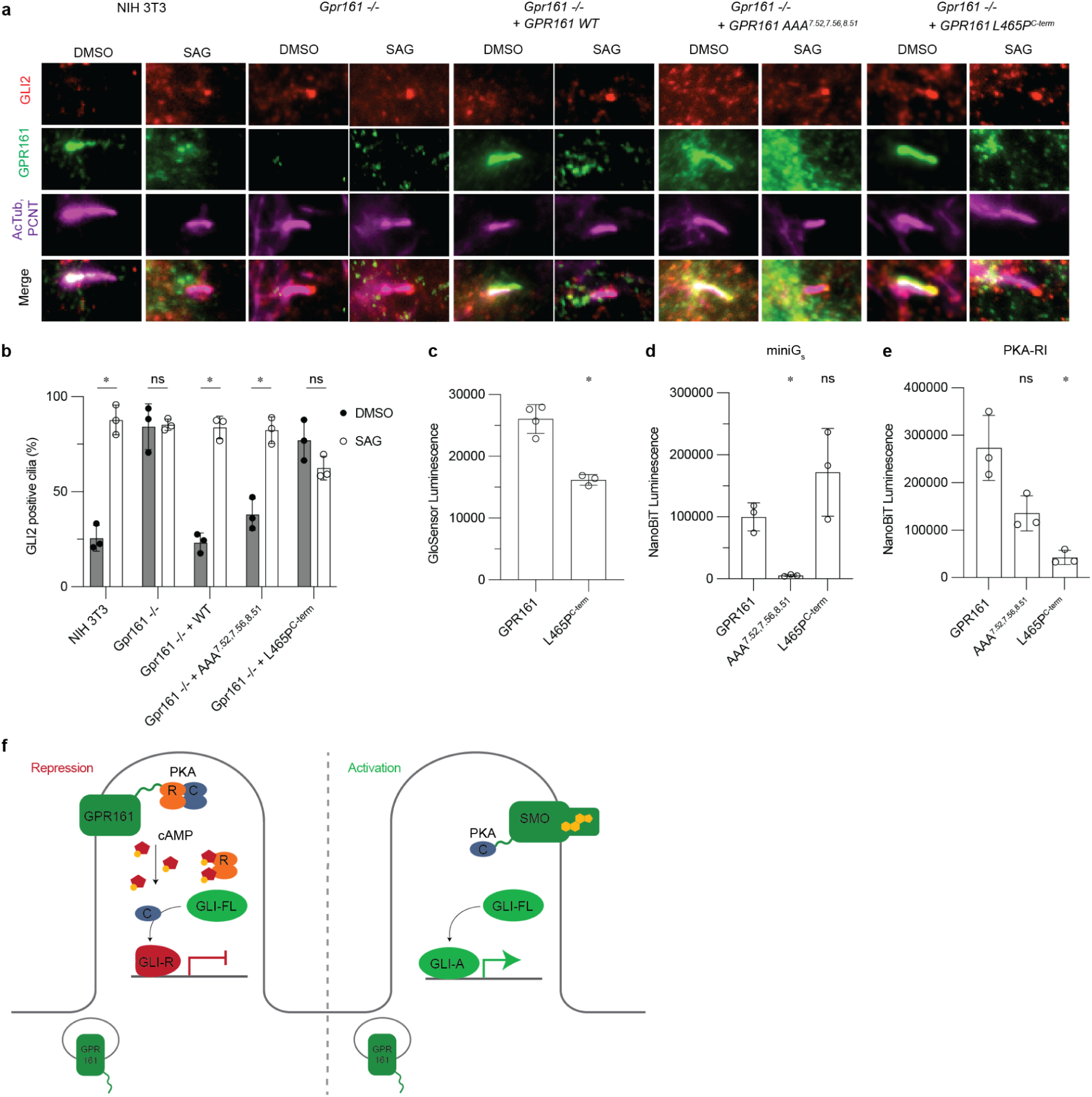
GPR161 PKA-RI binding, but not cAMP generation, is necessary to repress ciliary trafficking of GLI2. **a)** Representative images of the effect of site-directed mutagenesis of GPR161 on ciliary localization and GLI2 repression in ciliary tips in NIH 3T3 cells. NIH 3T3 Flp-In CRISPR based *Gpr161*^-/-^ cells stably expressing untagged mouse wild-type or *GPR161* mutants were starved for 24 hr upon confluence and were treated for further 24 hr ± SAG (500 nM). After fixation, cells were immunostained with anti-GLI2 (red), anti-GPR161 (green), anti-acetylated, and centrosome (AcTub; PCNT purple) antibodies. Scale bar, 5 µm. **b)** Quantification GLI2 positive cilia indicating Hedgehog pathway activation. AAA^7.52, 7.56, 8.51^ rescues function similar to WT, and L465P^C-term^ does not, similar to *Gpr161*^-/-^. Quantification of GLI2 positive cilia are shown from 3 biologically independent experiments from images taken from 2-3 different regions/experiment and counting 15-30 cells/region. Data are mean ± s.d. (\**P* < 0.05; ns, not significant; two-way ANOVA followed by Šidák’s multiple comparison tests). **c)** cAMP production assay assessing L465P^C-term^ mutation. Data are mean ± sd, n=3-4 biologically independent samples (\**P* < 0.05; one-way ANOVA followed by Dunnett’s multiple comparison tests). **d)** Nanoluc complementation assay for receptor recruitment of miniG_s_. Both GPR161 and L465P^C-term^ constitutively recruit miniG_s_ while AAA^7.52, 7.56, 8.51^ does not. Data are mean ± sd, n=3-4 biologically independent samples (\**P* < 0.05; ns, not significant; one-way ANOVA followed by Dunnett’s multiple comparison tests). **e)** Nanoluc complementation assay for receptor recruitment of PKA-RI. GPR161, AAA^7.52, 7.56, 8.51^, and L465P^C-term^ each recruit less PKA-RI, respectively. Data are mean ± sd, n=3-4 biologically independent samples (\**P* < 0.05; ns, not significant; one-way ANOVA followed by Dunnett’s multiple comparison tests). **f)** PKA-centric model of Hedgehog pathway repression in cilia. In the absence of Hedgehog, GPR161 represses the pathway in cilia through coupling PKA. GPR161 also functions in periciliary endosomal compartments in regulating GLI-R formation^6^. In the presence of Hedgehog, SMO activates the pathway by entering the cilia, binding cholesterol, and sequestering PKA-C, while GPR161 exits cilia.

### GPR161 induced cAMP is dispensable for repression of GLI2 ciliary accumulation

We next aimed to understand how activation of GPR161 leads to Hedgehog pathway repression in cilia. Upon Hedgehog pathway activation, GLI2 proteins accumulate at the tips of cilia^43–45^. We previously showed removal of GPR161 from cilia, either as a *Gpr161* gene knockout or from disruption of GPR161 trafficking to cilia results in accumulation of GLI2 in ciliary tips of resting cells^6^. We therefore used GLI2 ciliary accumulation as a sensitive test for GPR161 function in this cellular compartment.

GPR161 localizes to primary cilia in ciliated NIH 3T3 cells in the basal condition, assessed by co-localization with the ciliary markers acetylated tubulin (AcTub) and pericentrin (PCNT). Concordant with Hedgehog pathway inhibition, GLI2 does not accumulate in the primary cilium in the basal condition (DMSO treatment). Addition of the small molecule Hedgehog pathway agonist SAG leads to exit of GPR161 from cilia and accumulation of GLI2 in the ciliary tip (Fig. 5a). In *Gpr161^-/-^* NIH 3T3 cells, GLI2 accumulates in the ciliary tip in the basal condition and addition of SAG does not further increase ciliary GLI2 levels; this phenotype is consistent with loss of ciliary Hedgehog pathway repression by GPR161. Expression of wild-type GPR161 in *Gpr161^-/-^* NIH 3T3 cells rescues repression of Hedgehog pathway in cilia, as indicated by a low level of GLI2 positive cilia in the basal condition.

We first examined whether GPR161 ECL2 mutants, which are defective for cAMP production, can rescue GLI2 ciliary suppression. Here, we found that ECL2 mutants of GPR161 failed to accumulate in the primary cilium, suggesting that these mutants are either defective in biogenesis or are impaired in normal trafficking to the primary cilium (Supplementary Fig. 9b). Consistent with a lack of GPR161 localization in cilia, GPR161 ECL2 mutants also failed to suppress GLI2 in ciliary tips.

We next turned to the GPR161-AAA^7.52, 7.56, 8.51^ and GPR161-V129E^3.54^ mutants to understand whether cholesterol-dependent G_s_ activation is important for GPR161 repression of the Hedgehog pathway in cilia. Unlike the ECL2 mutants, GPR161-AAA^7.52, 7.56, 8.51^ and GPR161-V129E^3.54^ expressed in *Gpr161^-/-^* NIH 3T3 cells showed robust ciliary localization similar to wild-type GPR161 (Fig. 5a. and Supplementary Fig. 9a,b). Like wild-type GPR161, activation of the Hedgehog pathway by SAG led to exit of GPR161-AAA^7.52, 7.56, 8.51^ from the primary cilium. By contrast, GPR161-V129E^3.54^ did not exit cilia upon Hedgehog pathway activation (Supplementary Fig. 9a,b). Our prior studies with GPR161-V129E^3.54^ suggested that this mutant reduced cAMP production^1^; we found here that GPR161-V129E^3.54^ did not suppress ciliary GLI2 when expressed in *Gpr161^-/-^* NIH 3T3 cells (Supplementary Fig. 9a,c). However, the inability of GPR161-V129E^3.54^ to exit cilia, combined with residual interaction with miniG_s_, suggests that this mutation may have consequences beyond decreased cAMP production. Surprisingly, expression of the GPR161-AAA^7.52, 7.56, 8.51^ mutant suppressed GLI2 accumulation, consistent with repression of the Hedgehog pathway in the basal state (Fig. 5a,b). This unexpected result suggests that a G_s_ signaling-independent function of GPR161 is the predominant mediator for GLI repression in cilia in NIH 3T3 cells.

GPR161 is unique among many GPCRs in that it contains a PKA-binding AKAP domain. Previous studies have identified an amphipathic helix in the C-terminus of GPR161 that directly binds PKA regulatory subunits type I (RI)^25^; introduction of a single point mutant, L465P^C-term^, breaks this amphipathic helix and prevents PKA type I holoenzyme binding. Compared to the sterol-binding site, this PKA-binding site in GPR161 is less well conserved (Supplementary Fig. 6b). We assessed whether disruption of PKA binding by GPR161 influences Hedgehog pathway repression. In HEK293 cells, expression of GPR161-L465P^C-term^ led to constitutive cAMP production, albeit to a lesser extent than wild-type GPR161 (Fig. 5c). Indeed, GPR161-L465P^C-term^ robustly recruited miniG_s_ in a NanoLuciferase fragment complementation assay^33^ (Fig. 5d), indicating that this mutation does not influence GPR161 interactions with G_s_. Consistent with previous reports, GPR161-L465P^C-term^ recruited less PKA-RI than wild-type GPR161 (Fig. 5e). By contrast, the cAMP deficient mutants GPR161-AAA^7.52, 7.56, 8.51^ and GPR161-V129E^3.54^ recruited PKA-RI, albeit to slightly lower levels than the wild-type GPR161 (Fig. 5e and Supplementary Fig. 9e).

Having validated that GPR161-L465P^C-term^ attenuates interaction with PKA-RI, we tested whether this mutant represses the ciliary Hedgehog pathway in *Gpr161^-/-^* NIH 3T3 cells. We validated that GPR161-L465P^C-term^ was located in primary cilia in the basal condition (Fig. 5a). Like wild-type GPR161, GPR161-L465P^C-term^ exited cilia upon activation of the Hedgehog pathway by SAG (Fig. 5a and Supplementary Fig. 9b). However, GPR161-L465P^C-term^ was unable to repress GLI2 localization to the ciliary tip in the basal condition, indicating that PKA anchoring by GPR161 is critical for ciliary Hedgehog pathway control. Indeed, a double mutant combining disruption of G_s_ coupling and PKA binding (GPR161-AAA^7.52, 7.56, 8.51^-L465P^C-term^) was also unable to repress ciliary GLI2 in the basal condition (Supplementary Fig. 9c). We conclude that GPR161 binding to PKA-RI is essential for Hedgehog repression in the primary cilia, while GPR161-induced G_s_ signaling is dispensable.

## Discussion

Our studies here illuminate fundamental mechanisms of GPR161 activation, and how these mechanisms relate to GPR161-mediated regulation of Hedgehog signal transduction. Our cryo-EM structure of GPR161 revealed two stimuli contributing to GPR161 constitutive activity: first, a self-activating ECL2; and second, a sterol-like density at a unique extrahelical site. We demonstrate that ECL2 occludes the canonical Class A GPCR orthosteric site and is required for GPR161 trafficking to the primary cilia and cAMP signaling. We show that cholesterol can bind at the sterol site, that sterol-binding site availability is dependent upon the G protein-bound conformation of GPR161, and that the sterol-binding site regulates cAMP signaling. These two features of GPR161 activation illuminate the basis for G_s_-induced cAMP constitutive activity observed in previous studies^1,6,21^.

With these fundamental activation properties of GPR161, we provide new context into how GPR161 regulates the Hedgehog signaling pathway. A central model for GPR161 function in Hedgehog pathway repression is the importance of constitutive cAMP generation^1^. Optogenetic and chemogenetic triggers that elevate ciliary cAMP levels repress Hedgehog signaling^17^. We directly tested the importance of GPR161-induced cAMP production in one aspect of Hedgehog pathway repression, namely suppression of GLI2 transcription factor accumulation in the primary cilium in NIH 3T3 cells. We previously showed that a complete lack of GPR161 from cilia alone is important in suppressing GLI2 levels in the primary cilium^6^. Thus, this assay is a precise read out of GPR161 activity in cilia. Surprisingly, a GPR161 mutant that is unable to couple to G_s_ and support cAMP production (GPR161-AAA) retains the ability to suppress ciliary GLI2 accumulation in NIH 3T3 cells, suggesting that cAMP production of GPR161 is less crucial for Hedgehog pathway repression in cilia than current models suggest. Rather, we find that preventing the anchoring of the PKA type I complexes to GPR161 plays a more important role in suppressing GLI2 levels in cilia.

We propose the following model for repression of the Hedgehog pathway by GPR161 in cilia (Fig. 5f). In the absence of Hedgehog, GPR161 bound to PKA is localized to primary cilia. PKA within the cilia phosphorylates GLI resulting in processing into its repressor form. The general presence of ciliary cAMP is important for this process but could be generated by other ciliary GPCRs^23^, by receptor independent activation^46^ of adenylyl cyclases by G_s_ in cilia, or by G-protein independent activity of adenylyl cyclase^47^. A complete disruption of transport of adenylyl cyclases into cilia from upstream maturation defects during ER-Golgi transit in the secretory pathway, as seen in the *Ankmy2* knockout, prevents GLI-R formation in embryos and results in GLI2 accumulation in cilia^48^. In addition, although basal levels of cAMP in cilia are controversial, with reports ranging from levels comparable to that in the cytoplasm^49^ to supraphysiological levels ∼4.5 µM^47^, PKA-C can be activated with sub-optimal cAMP levels^50^. In the presence of Hedgehog, GPR161 traffics out of the cilia, removing PKA-RI with it^51^.

The above model does not exclude the role of extraciliary GPR161, particularly that in the periciliary endosomal compartment^6^, in GLI-R processing and thereby regulating tissue-specific morpho-phenotypes. We have recently demonstrated that GPR161 is not only localized to primary cilia but is also located in periciliary endosomes^6^. Both ciliary and extraciliary pools of GPR161 contribute to GLI-R formation and regulate tissue-specific repression of Hedgehog pathway in mice^6^. Although we show that GPR161 AKAP activity is critical for suppression of GLI2 trafficking in cilia, the AKAP function of GPR161 is not fully necessary for its suppression of Hedgehog pathway phenotypes in zebrafish^20^. The most parsimonious model explaining these paradoxical results would be that while ciliary AKAP function of GPR161 is critical for suppression of GLI2 trafficking to cilia, GPR161 functions outside cilia through sterol-mediated cAMP signaling. Production of cAMP has been reported for other GPCRs in the endosomal compartment^52–55^. Further experiments in organismal models will be needed to test the role of cAMP and AKAP signaling by GPR161 in the extraciliary endosomal compartment in tissue-specific Hedgehog signaling.

Our revised model for GPR161 provides a compelling parallel to recent reports that highlight direct interactions between Smoothened and PKA-C in Hedgehog signaling. As a G_i_-coupled GPCR, Smoothened suppression of cAMP generation was initially described as critical for Hedgehog signaling in the fly^56^. Subsequent studies in vertebrates have called into question the importance of Smoothened-induced G_i_ signaling in cilia in vertebrates^19,57^. More recently, the identification of a PKA-inhibitory motif in the Smoothened C-terminus suggests that activated Smoothened directly sequesters the catalytic subunits of PKA (PKA-C) to suppress enzymatic activity^58^. Instead of acting via cAMP on PKA, we propose that two GPCRs important to Hedgehog signaling, GPR161 and Smoothened, predominantly depend on binary interactions with PKA-C or RI subunit complexes to regulate the Hedgehog pathway in cilia (Fig. 5f).

Our identification of a conserved sterol binding site in GPR161 raises fundamental questions about the role of sterols in control of GPR161 signaling. We compared relative conservation of the GPR161 sterol binding site to the PKA-binding helix in the C-terminus (Supplementary Fig. 6b). Although the sterol binding site is conserved in deuterostome genomes, the PKA-binding motif is not clearly identified in echinoderms (e.g., *S. purpuratus*), early branching chordates (e.g., *B. belcheri*), and hemichordates (e.g., *S. kowalevskii*). The strong conservation of the cholesterol binding site, and the importance of this site for GPR161 to couple to G_s_ and generate cAMP, points to sterol driven cAMP generation of GPR161 having an important biological function.

Our finding that the conserved sterol binding site is not critical for controlling GLI2 levels in the primary cilium suggests several possibilities. First, as noted above, it is possible that extraciliary GPR161 uses a mechanism distinct from its ciliary AKAP function to control the Hedgehog pathway. For example, extraciliary GPR161 may depend on cholesterol or other sterols to promote cAMP formation to control GLI-R formation. Second, it is entirely possible that cAMP production by GPR161 has roles in adult tissues outside of the Hedgehog pathway. Indeed, GPR161 is expressed in many different cell types in adult tissues that are not hedgehog regulated, such as the adult hippocampus CA1 region^59^. GPR161 has also been reported to localize to cilia of hippocampal neurons^1,59,60^. Ciliary peptidergic GPCR signaling in the CA1 pyramidal neurons has been recently shown to regulate chromatin accessibility^61^, but the role of cAMP signaling mediated by cilia remains unknown. Third, GPR161 could also have roles in cancers beyond its role in Hedgehog pathway repression in medulloblastoma^3,7^. For example, *GPR161* is overexpressed in triple-negative breast cancer and has been proposed to promote cell proliferation and invasiveness in tumor cells by forming a signaling complex with β-arrestin2 and IQGAP1^4,62^. Our identification of a GPR161 mutant that specifically attenuates cAMP production will enable a careful dissection of these potential roles of GPR161, irrespective of its function in cilia and beyond the role proposed in the Hedgehog pathway.

Our study highlights how GPCR cAMP-PKA signaling establishes precise signaling microdomains in primary cilia. Such restrictive signaling in nanodomains has been an emerging feature of subcellular signaling by cAMP^63,64^. Most broadly, our work highlights that orphan GPCRs may have functions beyond the biological pathway where they are first encountered. Directly observing the stimuli that activate orphan GPCRs will enable precise approaches to dissect the functional relevance of a specific signaling pathway in a biological outcome. The advent of structure-based methods to interrogate orphan GPCRs will therefore broaden views on the possible biology coordinated by this fascinating family of understudied proteins.

## Methods

### Expression and purification of GPR161 active-state complex

The human *GPR161* gene with an N-terminal influenza hemagglutinin signal sequence and FLAG epitope was cloned into a pcDNA3.1 vector with zeocin resistance and a tetracycline inducible cassette, as described previously^66^. The miniG_s_399 protein^26^ was fused to the C terminus of GPR161 preceded by a glycine/serine linker and rhinovirus 3C protease recognition site. The resulting fusion construct was transfected into inducible Expi293F-TetR cells (unauthenticated and untested for mycoplasma contamination, Thermo Fisher) using the ExpiFectamine transfection reagent per manufacturer instructions. After 18 h, protein expression was induced with 1 µg/mL doxycycline hyclate for 28 h before collection by centrifugation. Pelleted cells were washed with 50 mL phosphate buffered saline, pH 7.5 before storage at −80 °C.

For complex purification for cryo-EM, frozen cells were hypotonically lysed in a buffer comprised of 20 mM HEPES pH 7.5, 1 mM EDTA, 160 µg/mL benzamidine, 160 µg/mL leupeptin, and 100 µM TCEP for 10 min at 25°C. The membrane fraction was collected by centrifugation, and the fusion protein was extracted with a buffer comprised of 20 mM HEPES, pH 7.5, 300 mM NaCl, 1% (w/v) lauryl maltose neopentyl glycol (L-MNG, Anatrace), 0.1% (w/v) cholesteryl hemisuccinate (CHS, Steraloids), 2 mM MgCl_2_, 2 mM CaCl_2_, 160 µg/mL benzamidine, 2 µg/mL leupeptin, and 100 µM TCEP with dounce homogenization and incubation with stirring for one hour at 4 °C. The soluble fraction was separated from the insoluble fraction by centrifugation and was applied to a column of homemade M1–FLAG antibody-conjugated Sepharose beads at a rate of 1 mL/min. Sepharose resin was then washed with ten column volumes of 20 mM HEPES pH 7.5, 300 mM NaCl, 0.1% (w/v) L-MNG, 0.01% (w/v) CHS, 2 mM MgCl_2_, 2 mM CaCl_2_, 100 µM TCEP and then washed with 10 column volumes of 20 mM HEPES pH 7.5, 150 mM NaCl, 0.0075% (w/v) L-MNG, 0.00075% (w/v) CHS, 2 mM MgCl_2_, 2 mM CaCl_2_, 100 µM TCEP prior to elution with 20 mM HEPES pH 7.5, 150 mM NaCl, 0.0075% (w/v) L-MNG, 0.00075% (w/v) CHS, 100 µM TCEP, 5 mM EDTA, and 0.2 mg/mL FLAG peptide. The eluted GPR161-miniG_s_ fusion protein was concentrated in a 100 kDa MWCO Amicon spin concentrator, and injected onto a Superdex 200 Increase 10/300GL (Cytiva) gel filtration column equilibrated in 20 mM HEPES pH 7.5, 150 mM NaCl, 100 µM TCEP, 0.0075% (w/v) L-MNG, 0.0025% glyco-diosgenin (GDN, Anatrace), and 0.0005% CHS. Monodisperse fractions of GPR161-miniG_s_ were complexed with G_β1γ2_ heterodimer and Nb35 (purified as described previously ^67^) at 2-fold molar excess overnight at 4°C. The next day, the complex was concentrated with a 100 kDa MWCO spin concentrator and excess G_β1γ2_ and Nb35 was removed via size-exclusion chromatography, using a Superdex200 Increase 10/300 GL column (GE Healthcare) equilibrated in 20 mM HEPES pH 7.5, 150 mM NaCl, 100 µM TCEP, 0.0075% (w/v) L-MNG, 0.0025% (w/v) GDN, and 0.00075% CHS. The resulting complex was concentrated to 2.9 mg/mL with a 100 kDa MWCO spin concentrator for preparation of cryo-EM grids.

Two separate preparations of GPR161-miniG_s_ were made for biochemical experiments that deviated slightly from the purification protocol for cryoEM. For cholesterol photolabeling experiments, GPR161-miniG_s_ was expressed and purified using the above protocol except CHS and GDN was excluded from all buffers. For ^3^H-cholesterol binding experiments, n-Dodecyl-*β*-D-Maltopyranoside (DDM, Anatrace) was used in lieu of LMNG at a final concentration of 0.02% and CHS and GDN were excluded from all buffers. The resulting size exclusion chromatography-purified protein samples were flash frozen in liquid nitrogen for downstream assay use.

### Cryo-EM vitrification, data collection, and processing

The GPR161-G_s_-Nb35 complex was concentrated to 3 mg/mL and 3 µl was applied onto a glow-discharged 300 mesh 1.2/1.3 gold grid covered in a holey carbon film (Quantifoil). Excess sample was removed with a blotting time of 3 s and a blotting force of 1 at 4 °C prior to plunge freezing into liquid ethane using a Vitrobot Mark IV (Thermo Fisher). A total of 8,294 movies were recorded with a K3 detector (Gatan) on a Titan Krios (Thermo Fisher) microscope operated at 300 keV with a BioQuantum post-column energy filter set to a zero-loss energy selection slit width set of 20 eV. The 60-frame movies were recorded for 2.6 s at a physical pixel size of 0.86 Å per pixel and a defocus range of −0.8 to − 2.2 µm for a total dose of 50.7 e−/Å2. Exposure areas were acquired with an automated image shift collection strategy using EPU (Thermo Fisher).

Movies were motion-corrected and dose-fractionated on-the-fly during data collection using UCSF MotionCor2^68^. Corrected micrographs were imported into cryoSPARC v3.1 (Structura Biotechnology)^69^ for CTF estimation via the Patch CTF Estimation job type. Micrographs with a CTF fit resolution of > 5 Å were excluded from further processing. Templates for particle picking were generated using a 20 Å low-pass filtered model generated from an *ab initio* model made from blob-picked and 2D classified particles. Template picking yielded 9,760,777 particles, which were extracted in a 288-pixel box and Fourier cropped to 72 pixels. Particles were classified in 3D with alignment using the 20 Å low-pass filtered *ab initio* model and three “random” reconstructions generated from a prematurely terminated *ab initio* reconstruction job, called “garbage collectors,” with the Heterogeneous Refinement job type. Two rounds of Heterogeneous Refinement yielded 3,033,326 particles that were re-extracted in the same box sized cropped to 144 pixels. Additional Heterogeneous refinement and extraction without binning yielded 1,143,566 particles that were refined using the Non-Uniform Refinement job type. Particles were exported using csparc2star.py from the pyem script package^70^, and a mask covering the 7TM domain of GPR161 was generated using the Segger tool in UCSF ChimeraX^71^ and the mask.py script in pyem. The particles and mask were imported into Relion v3.0^72^ and classified in 3D without alignment. Particles comprising the three highest resolution classes were reimported into cryoSPARC for Non-Uniform Refinement. Finally, particles were exported into cisTEM^73^ for two local refinements using the Manual Refinement job type and low-pass filtering outside of masks. In the first local refinement, the previous 7TM mask was used, and the second local refinement used a full-particle mask.

### Model building and refinement

Model building and refinement began with the Alphafold2^74^ predicted structure as the starting model, which was fitted into the experimental cryoEM map using UCSF ChimeraX. The model was iteratively refined with real space refinement in Phenix^75^ and manually in Coot^76^. The cholesterol model and rotamer library were generated with the PRODRG server^77^, docked using Coot, and refined in Phenix. Final map-model validations were carried out using Molprobity and EMRinger in Phenix.

### cAMP signaling assays

We measured cAMP production to determine activation of G_s_ signaling by GPR161. For each GPR161 construct (wild-type (WT), M177R, V179R, W182G, W182R, W327A, W327R, R332A, AA, AAA, L465P), a 2 mL suspension culture of Expi293F-TetR cells was co-transfected with a 3:1 ratioof a pcDNA3.1 plasmid expressing GPR161 and a luciferase-based cAMP biosensor, pGlosensor-22F (Promega). Surface expression levels of constructs were titrated to similar levels with doxycycline and measured by flow cytometry using an Alexa-647 conjugated anti-M1 Flag antibody. Cells were collected 24 h post-induction, resuspended in Expi293 expression media (Gibco) supplemented with 10% DMSO, and gradually frozen to -80 °C in a Mr. Frosty Freezing container for future use. To perform the assay, frozen cells were rapidly thawed in a 37°C water bath and resuspended in fresh Expi293 expression medium. Cells were diluted to a final concentration of 1e6 cells per mL in Expi293 expression medium plus 2% (v/v) Glosensor assay reagent (Promega) and intubated for 75 min at room temperature with gentle rotation. Cells were then plated in a white 384-well plate (Greiner) to a final density of 15,000 cells per well. Immediately after cell addition, luminescence was measured using a CLARIOstar instrument. Statistical analyses were performed using one-way ANOVA followed by Dunnett’s multiple comparison tests between all possible pairs using GraphPad Prism 9 (Dotmatics).

### 3H-Cholesterol binding assay

To measure cholesterol binding to GPR161, we developed a scintillation proximity assay (SPA) using purified receptor and ^3^H-cholesterol (PerkinElmer). To capture M1-FLAG tagged receptor, we used Protein A coated beads and purified M1-FLAG antibody. Each binding reaction was performed in a final volume of 100 µL in a binding buffer comprised of 0.01% dodecylmaltoside, 20 mM HEPES pH 7.5, 150 mM NaCl, 100 nM purified M1-FLAG antibody, and 2 mM CaCl_2_ (to enable M1-FLAG tag binding to antibody). SPA beads were added to a final concentration of 0.675 mg/mL, ^3^H-cholesterol was added to 100 nM, and 100 nM of purified GPR161-miniG_s_ or GPR161-AAA^7.52, 7.56, 8.51^-miniG_s_ was added to start the reaction. For competition with cold cholesterol a 3 µM final concentration was used. The reactions were incubated with shaking at room temperature for 24 hrs, and bound ^3^H-cholesterol was measured in a scintillation counter (Perkin Elmer). Statistical analyses were performed using two-way ANOVA followed by Dunnett’s multiple comparison tests between all possible pairs using GraphPad Prism 9 (Dotmatics).

### Photolabeling and MS analysis

Photolabeling reagents were synthesized as previously described^39–41^ and stored in the dark at -20°C as 10 mM stocks in ethanol. Aliquots of the photolabeling reagents (KK231 or LKM38) were air dried in the dark at room temperature and resolubilized in 20 mM HEPES pH 7.5, 150 mM NaCl, 0.0075% LMNG (no CHS or GDN) containing 20 µg of purified GPR161-miniG_s_ in a total volume of 50 µL. The protein was incubated with the photolabeling regent for one hour in the dark at 4 °C and then irradiated in a quartz cuvette with >320 nm UV light as previously described^78^. For site identification experiments, the photolabeling reagent concentration was 100 µM. For cholesterol competition experiments, aliquots of ethanolic stocks of the photolabeling reagent and cholesterol (10 mM stock) were added into the same tube and air dried prior to solubilization with GPR161-miniG_s_. The final concentration of the photolabeling reagents was 3 µM and of cholesterol 100 µM.

For mass spectrometric analysis, the samples were desalted using Biospin 6 columns (BioRad, CA), equilibrated with 50 mM triethylammonium bicarbonate and 0.02%(w/v) n-Dodecyl-β-D-Maltoside. The proteins were sequentially reduced with 5 mM tris(2-carboxyethyl)phosphine (TCEP) for 30 mins, alkylated with 5 mM N-ethylmaleimide (NEM) in the dark for 45 mins, and quenched with 5 mM dithiothreitol (DTT) for 15 mins. These three steps were done at room temperature. The proteins were digested with 8 µg trypsin at 4 °C for one week at which time the digestions were terminated by addition of formic acid (FA) to a final concentration of 1%.

The resultant peptides were analyzed with an OrbiTrap ELITE mass spectrometer (Thermo Fisher Scientific). Briefly, 15 μL samples were injected by an autosampler (UltiMate 3000 UHPLC system; ThermoFisher) onto a home-packed polymeric reverse phase PLRP-S (Agilent, Santa Clara, CA) column (10 cm × 75 μm, 300 Å) at a flow rate of 800 nL/min. A 10%-95% acetonitrile (ACN) gradient was applied for 150 minutes to separate peptides. Solvent A was 0.1% FA/water, and solvent B was 0.1% FA/ACN. The following gradient was applied: isocratic elution at 10% solvent B, 1–60 minutes; 10%–95% solvent B, 60–110 minutes; 95% solvent B, 110–140 minutes; 95%–10% solvent B, 140–145 minutes; 10% solvent B, 145–150 minutes. For the first 60 minutes, a built-in divert valve on the mass spectrometer was used to remove the hydrophilic contaminants from the mass spectrometer. Mass spectra (MS1) were acquired at high resolution (resolution of 60,000) in the range of m/z = 100-2,000. Top 20 ion precursors in MS1 were selected for MS2 using data-dependent acquisition with exclusion of singly charged precursors. Fragmentation was performed with high-energy dissociation (HCD) using a normalized collision energy of 35%. Product ion spectra (MS2) were acquired at a resolution of 15,000.

The data were searched against a database containing the sequence of GPR161-miniG_s_ using PEAKS Studio X pro (Bioinformatics Solutions Inc, Waterloo, ON, Canada) with the following settings: precursor ions mass accuracy of 20 ppm, fragmentation ion accuracy of 0.1 Da, up to three missed cleavages on either side of peptide with trypsin digestion; methionine oxidation, cysteine alkylation with NEM and DTT, any amino acids with adduct of LKM38 (mass = 396.34) and KK231 (mass = 484.26) were included as variable modifications. The searched results were filtered with a 1% false discovery rate and the detected peptides were confirmed by manual analysis for monoisotopic mass accuracy and retention time with Xcalibur 2.2 (ThermoFisher). Fragment ions were also manually confirmed and were accepted based on the presence of a monoisotopic mass within 20 ppm mass accuracy. Photolabeling efficiency was estimated by generating selected ion chromatograms (SIC) of both unlabeled and photolabeled peptides, determining the area under the curve and calculating efficiency as: labeled peptide / (unlabeled peptide + labeled peptide). Statistical significance was analyzed with Student’s paired t-test using GraphPad Prism 9 (Dotmatics).

### NanoBiT recruitment assays

We measured protein recruitment to determine functionality of GPR161 mutants. For each GPR161 construct (wild-type (WT), AAA, L465P, V129E), as well as for β_2_AR and GPR52, a 2-mL suspension culture of Expi293F-TetR cells was co-transfected with a 500 ng of a pcDNA3.1 plasmid expressing receptor fused C-terminally to smBiT and 100 ng of transducer fused to lgBiT (miniG_s_: N-terminally, PKA-RI: C-terminally). Surface expression levels of constructs were titrated to similar levels with doxycycline and measured by flow cytometry using an Alexa-647 conjugated anti-M1 FLAG antibody. After 24 h of induction, cells were centrifuged at 300x*g*, and resuspended in DPBS at a concentration of ∼55,000 cells per 200 µL. 40 µL of 30 µM coelenterazine-h diluted in PBS was added to cells for a final concentration of 5 µM. Cells were incubated for ∼30 min at room temperature with gentle shaking. Luminescence was measured using a CLARIOstar instrument. Statistical analyses were performed using one-way ANOVA followed by Dunnett’s multiple comparison tests between all possible pairs using GraphPad Prism 9 (Dotmatics).

### Molecular dynamics simulations

#### Simulation setup

We performed simulations of GPR161 with cholesterol under two conditions: A) GPR161 unrestrained (6 independent simulations, roughly 1 μs each) B) GPR161 restrained to its G protein—bound conformation (6 independent simulations, roughly 1 μs each). For all conditions, the initial structures were based on the cryoEM structure reported [MOU1] in this paper and were prepared using Maestro (Schrödinger, LLC). In both conditions, the nanobody and G protein were removed from the structure.

Missing amino acid side chains were modeled using Prime (Schrödinger, LLC). Neutral acetyl and methylamide groups were added to cap the N- and C-termini, respectively, of the protein chains. Titratable residues were kept in their dominant protonation state at pH 7.4. Histidine residues were modeled as neutral, with a hydrogen atom bound to either the delta or epsilon nitrogen depending on which tautomeric state optimized the local hydrogen-bonding network. Dowser ^79^ was used to add water molecules to protein cavities. GPR161 was aligned on the receptor in the crystal structure of Prostaglandin E2 receptor EP2 subtype (PDB ID: 7CX4) ^80^ in the Orientation of Proteins in Membranes (OPM) database ^81^. The aligned structures were inserted into a pre-equilibrated palmitoyloleoyl-phosphatidylcholine (POPC) membrane bilayer using Dabble ^82^. Sodium and chloride ions were added to neutralize each system at a concentration of 150 mM. To simulate the G protein–bound conformation in condition B, 0.5 kcal·mol^-1^ ·Å^-2^ restraints were applied throughout the production simulation on non-hydrogen atoms of GPR161 residues that are within 4 Å of the G protein in the experimentally determined structure. These residues were: 125, 128, 129, 130, 131, 132, 133, 135, 136, 137, 211, 214, 215, 218, 219, 267, 268, 271, 272, and 327. Both final systems consist of 52716 atoms, including 140 lipid molecules and 9810 water molecules (initial system dimensions: 85 Å x 80 Å x 82 Å).

#### Simulation protocols

For each simulation, initial atom velocities were assigned randomly and independently. We employed the CHARMM36m force field for protein molecules, the CHARMM36 parameter set for lipid molecules and salt ions, and the associated CHARMM TIP3P model for water ^83,84^. Simulations were run using the AMBER20 software ^85^ under periodic boundary conditions with the Compute Unified Device Architecture (CUDA) version of Particle-Mesh Ewald Molecular Dynamics (PMEMD) ^86^ on one GPU.

The systems were first heated over 12.5 ps from 0 K to 100 K in the NVT ensemble using a Langevin thermostat with harmonic restraints of 10.0 kcal·mol^-1^ ·Å^-2^ on the non-hydrogen atoms of the lipids, protein, and cholesterol. Initial velocities were sampled from a Boltzmann distribution. The systems were then heated to 310 K over 125 ps in the NPT ensemble. Equilibration was performed at 310 K and 1 bar in the NPT ensemble, with harmonic restraints on the protein and cholesterol non-hydrogen atoms tapered off by 1.0 kcal·mol^-1^ ·Å^-2^ starting at 5.0 kcal·mol^-1^ ·Å^-2^ in a stepwise manner every 2 ns for 10 ns, and finally by 0.1 kcal·mol^-1^ ·Å^-2^ every 2 ns for an additional 18 ns. Except for the restrained residues listed above in condition B, all restraints were completely removed during production simulation. Production simulations were performed at 310 K and 1 bar in the NPT ensemble using the Langevin thermostat and Berendsen barostat.

Lengths of bonds to hydrogen atoms were constrained using SHAKE, and the simulations were performed using a timestep of 4.0 fs while using hydrogen mass repartitioning ^87^. Non-bonded interactions were cut off at 9.0 Å, and long-range electrostatic interactions were calculated using the particle-mesh Ewald (PME) method with an Ewald coefficient (β) of approximately 0.31 Å and B-spline interpolation of order 4. The PME grid size was chosen such that the width of a grid cell was approximately 1 Å. Snapshots of the trajectory were saved every 200 ps.

#### Simulation analysis protocols

The AmberTools17 CPPTRAJ package ^88^ was used to reimage trajectories at 1 ns per frame, Visual Molecular Dynamics (VMD) ^89^ was used for visualization and analysis, and PyMOL (The PyMOL Molecular Graphics System, Schrödinger, LLC) was used for renderings.

Plots of time traces from representative simulations were generated with Matplotlib ^90^ and show both original, unsmoothed traces (transparent lines) and traces smoothed with a moving average (thick lines), using an averaging window of 20 ns. All traces include the initial equilibration with harmonic restraints on the protein and cholesterol non-hydrogen atoms.

To monitor ECL2 movement, we measure the minimal distance between all atoms of W182 and T189 (Fig. 2f,g and Supplementary Fig. 8b). To capture cholesterol motion, we measure the minimal distance between all non-hydrogen atoms between cholesterol and W327. We show a representative trace of this measurement for each condition (Fig. 4b and Supplementary Fig. 8c,d).

### Ciliary localization and Hedgehog pathway activation

#### Cell lines

NIH 3T3-FlpIn cells were authenticated by and purchased from Thermo Fisher Scientific. They have tested negative for Mycoplasma. The *Gpr161^-/-^*NIH 3T3 Flp-In cell line was a gift from Rajat Rohatgi^23^. The cells were cultured in DMEM-high glucose media (D5796; Sigma) with 10% BCS (Sigma-Aldrich), 0.05 mg/ml penicillin, 0.05 mg/ml streptomycin, and 4.5 mM glutamine. Stable knockout cell lines were generated by retroviral infection with pBABE constructs having untagged wild type or mutant *GPR161* inserts followed by antibiotic selection. Single or multiple amino acid mutations in full-length *GPR161* were generated using Q5 site-directed mutagenesis kit (NEB).

#### Immunostaining and microscopy

For immunofluorescence experiments in cell lines, cells were cultured on coverslips until confluent and starved for 48 h. To quantify ciliary GLI2 and GPR161 levels, cells were treated with 500 nM SAG or DMSO for 24 h after 24 h of serum starvation. Cells were fixed with 4% PFA for 10 min at room temperature. After blocking with 5% normal donkey serum, the cells were incubated with primary antibody solutions for 1 h at room temperature followed by treatment with secondary antibodies for 30 min along with DAPI. Primary antibodies used were against GPR161 (1:200, custom-made)^21^, acetylated α-tubulin (mAb 6-11B-1, Sigma; 1:2000), GLI2 (1:500, gift from Jonathan Eggenschwiler)^91^, pericentrin (611814, BD Biosciences; 1:500). Coverslips were mounted with Fluoromount-G and images were acquired with a Zeiss AxioImager.Z1 microscope using a 40x oil immersion objective lens.

#### Quantification and statistical analysis

Cilia positive for GLI2 or GPR161 in *Gpr161^-/-^* cells expressing untagged wild-type or mutant *GPR161* were counted. Statistical analyses were performed using two-way ANOVA followed by Šidák’s multiple comparison tests between all possible pairs using GraphPad Prism 9 (Dotmatics).

### Data Availability

Coordinates for GPR161-G_s_ complex have been deposited in the RCSB PDB under accession code 8SMV. EM density maps for GPR161-G_s_ have been deposited in the Electron Microscopy Data Bank under accession code EMD-40603. The MD simulation trajectories have been deposited in the Zenodo database under doi https://doi.org/10.5281/zenodo.7887650.

## Acknowledgments

This work was supported by the National Institutes of Health (NIH) grants R01GM127359 (R.O.D.), R01GM108799 (A.S.E.), P50MH122379 (A.S.E.), R35GM144136 (S.M.) and R01GM138992 (R.O.D. and A.M.). Additional support was from NSF Graduate Research Fellowship (M.K.) and Human Frontier Science Program Long-Term Fellowship LT000916/2018-L (C.-M.S.). Cryo-EM equipment at UCSF is partially supported by NIH grants S10OD020054 and S10OD021741. Some of this work was performed at the Stanford-SLAC Cryo-EM Center (S2C2), which is supported by the National Institutes of Health Common Fund Transformative High-Resolution Cryo-Electron Microscopy program (U24 GM129541). The authors would also like to thank the following S2C2 personnel for their invaluable support and assistance: Corey Hecksel. The authors thank Jeremy Reiter for helpful suggestions on GPR161 homologous sequences. The content is solely the responsibility of the authors and does not necessarily represent the official views of the National Institutes of Health. A.M. acknowledges support from the Edward Mallinckrodt, Jr. Foundation and the Vallee Foundation. A.M. is a Chan Zuckerberg Biohub Investigator.

## Contributions

N.H., S.H., and I.D. cloned, expressed, and biochemically optimized the purification of GPR161 complex constructs for structural studies. N.H., S.H., and I.D. performed cryo-EM data collection, with help from SLAC Cryo-EM Center, and data processing. N.H., S.H., I.D., and A.M. built and refined models of GPR161. N.H. and S.H. generated receptor constructs and determined expression levels by flow cytometry and performed signaling studies, complementation assays, and analyzed the data. M.K. and C.M.S. performed and analyzed molecular dynamics simulations under the supervision of R.O.D. N.H. prepared samples for, performed, and analyzed scintillation proximity assay data with A.M. Z.C. performed and analyzed mass spectrometry experiments using reagents provided by D.C. under the supervision of A.E. S.H.H., V.R.P, and S.M. prepared constructs, performed, and analyzed GPR161 localization and Hedgehog pathway repression experiments. S.B. performed phylogenetic analysis under the supervision of D.M. P.T., D.R., and E.S. analyzed GPR161 variants. All authors contributed to figures. N.H., S.H., A.M., and S.M. wrote the manuscript, with edits and approval from all authors. A.M. supervised the overall project.

## REFERENCES

1. Mukhopadhyay, S. et al. The Ciliary G-Protein-Coupled Receptor Gpr161 Negatively Regulates the Sonic Hedgehog Pathway via cAMP Signaling. Cell 152, 210–223 (2013).

2. Kim, S.-E. et al. Dominant negative GPR161 rare variants are risk factors of human spina bifida. Hum. Mol. Genet. 28, 200–208 (2019).

3. Begemann, M. et al. Germline GPR161 Mutations Predispose to Pediatric Medulloblastoma. J. Clin. Oncol. 38, 43–50 (2020).

4. Feigin, M. E., Xue, B., Hammell, M. C. & Muthuswamy, S. K. G-protein–coupled receptor GPR161 is overexpressed in breast cancer and is a promoter of cell proliferation and invasion. Proceedings of the National Academy of Sciences 111, 4191–4196 (2014).

5. Civelli, O., Saito, Y., Wang, Z., Nothacker, H.-P. & Reinscheid, R. K. Orphan GPCRs and their ligands. Pharmacol. Ther. 110, 525–532 (2006).

6. Hwang, S.-H., Somatilaka, B. N., White, K. & Mukhopadhyay, S. Ciliary and extraciliary Gpr161 pools repress hedgehog signaling in a tissue-specific manner. Elife 10, (2021).

7. Shimada, I. S. et al. Basal Suppression of the Sonic Hedgehog Pathway by the G-Protein-Coupled Receptor Gpr161 Restricts Medulloblastoma Pathogenesis. Cell Rep. 22, 1169–1184 (2018).

8. Hwang, S.-H. et al. The G protein-coupled receptor Gpr161 regulates forelimb formation, limb patterning and skeletal morphogenesis in a primary cilium-dependent manner. Development 145, (2018).

9. Kim, S.-E. et al. Wnt1 Lineage Specific Deletion of Gpr161 Results in Embryonic Midbrain Malformation and Failure of Craniofacial Skeletal Development. Front. Genet. 12, 761418 (2021).

10. Shimada, I. S. et al. Derepression of sonic hedgehog signaling upon Gpr161 deletion unravels forebrain and ventricular abnormalities. Dev. Biol. 450, 47–62 (2019).

11. Li, B. I. et al. The orphan GPCR, Gpr161, regulates the retinoic acid and canonical Wnt pathways during neurulation. Dev. Biol. 402, 17–31 (2015).

12. Matteson, P. G. et al. The orphan G protein-coupled receptor, Gpr161, encodes the vacuolated lens locus and controls neurulation and lens development. Proceedings of the National Academy of Sciences 105, 2088–2093 (2008).

13. Karaca, E. et al. Whole-exome sequencing identifies homozygous GPR161 mutation in a family with pituitary stalk interruption syndrome. J. Clin. Endocrinol. Metab. 100, (2015).

14. Website. https://bpspubs.onlinelibrary.wiley.com/doi/10.1111/bph.16053.

15. McMahon, A. P., Ingham, P. W. & Tabin, C. J. 1 Developmental roles and clinical significance of Hedgehog signaling. in Current Topics in Developmental Biology vol. 53 1–114 (Academic Press, 2003).

16. Kopinke, D., Norris, A. M. & Mukhopadhyay, S. Developmental and regenerative paradigms of cilia regulated hedgehog signaling. Semin. Cell Dev. Biol. 110, 89–103 (2021).

17. Truong, M. E. et al. Vertebrate cells differentially interpret ciliary and extraciliary cAMP. Cell 184, 2911–2926.e18 (2021).

18. Hilgendorf, K. I., Johnson, C. T. & Jackson, P. K. The primary cilium as a cellular receiver: organizing ciliary GPCR signaling. Curr. Opin. Cell Biol. 39, 84–92 (2016).

19. Mukhopadhyay, S. & Rohatgi, R. G-protein-coupled receptors, Hedgehog signaling and primary cilia. Semin. Cell Dev. Biol. 33, (2014).

20. Tschaikner, P. M. et al. Feedback control of the Gpr161-Gαs-PKA axis contributes to basal Hedgehog repression in zebrafish. Development 148, dev192443 (2021).

21. Pal, K. et al. Smoothened determines β-arrestin–mediated removal of the G protein–coupled receptor Gpr161 from the primary cilium. J. Cell Biol. 212, 861–875 (2016).

22. Kroeze, W. K. et al. PRESTO-Tango as an open-source resource for interrogation of the druggable human GPCRome. Nat. Struct. Mol. Biol. 22, 362–369 (2015).

23. Pusapati, G. V. et al. G protein–coupled receptors control the sensitivity of cells to the morphogen Sonic Hedgehog. Sci. Signal. 11, eaao5749 (2018).

24. Foster, S. R. et al. Discovery of Human Signaling Systems: Pairing Peptides to G Protein-Coupled Receptors. Cell 179, (2019).

25. Bachmann, V. A. et al. Gpr161 anchoring of PKA consolidates GPCR and cAMP signaling. Proc. Natl. Acad. Sci. U. S. A. 113, 7786–7791 (2016).

26. Nehmé, R. et al. Mini-G proteins: Novel tools for studying GPCRs in their active conformation. PLoS One 12, e0175642 (2017).

27. Carpenter, B. & Tate, C. G. Engineering a minimal G protein to facilitate crystallisation of G protein-coupled receptors in their active conformation. Protein Eng. Des. Sel. 29, 583–594 (2016).

28. Rasmussen, S. G. F. et al. Crystal structure of the β2 adrenergic receptor–Gs protein complex. Nature 477, 549–555 (2011).

29. Zhou, Q. et al. Common activation mechanism of class A GPCRs. Elife 8, (2019).

30. Lin, X. et al. Structural basis of ligand recognition and self-activation of orphan GPR52. Nature 579, 152–157 (2020).

31. Ye, F., et al. Cryo-EM structure of G-protein-coupled receptor GPR17 in complex with inhibitory G protein. MedComm (2020) 3, e159 (2022).

32. Wong, T.-S., et al. Cryo-EM structure of orphan G protein-coupled receptor GPR21. MedComm (2020) 4, e205 (2023).

33. Wan, Q. et al. Mini G protein probes for active G protein-coupled receptors (GPCRs) in live cells. J. Biol. Chem. 293, 7466–7473 (2018).

34. Dixon, A. S. et al. NanoLuc Complementation Reporter Optimized for Accurate Measurement of Protein Interactions in Cells. ACS Chem. Biol. 11, 400–408 (2016).

35. Copp, A. J. et al. Spina bifida. Nat Rev Dis Primers 1, 1–18 (2015).

36. Eaton, S. Multiple roles for lipids in the Hedgehog signalling pathway. Nat. Rev. Mol. Cell Biol. 9, 437–445 (2008).

37. Luchetti, G. et al. Cholesterol activates the G-protein coupled receptor Smoothened to promote Hedgehog signaling. Elife 5, (2016).

38. Cooper, M. K. et al. A defective response to Hedgehog signaling in disorders of cholesterol biosynthesis. Nat. Genet. 33, 508–513 (2003).

39. Budelier, M. M. et al. Photoaffinity labeling with cholesterol analogues precisely maps a cholesterol-binding site in voltage-dependent anion channel-1. J. Biol. Chem. 292, 9294–9304 (2017).

40. Krishnan, K. et al. Validation of Trifluoromethylphenyl Diazirine Cholesterol Analogues As Cholesterol Mimetics and Photolabeling Reagents. ACS Chem. Biol. 16, 1493–1507 (2021).

41. Castellano, B. M. et al. Lysosomal cholesterol activates mTORC1 via an SLC38A9-Niemann-Pick C1 signaling complex. Science 355, 1306–1311 (2017).

42. Shin, H. R. et al. Lysosomal GPCR-like protein LYCHOS signals cholesterol sufficiency to mTORC1. Science 377, 1290–1298 (2022).

43. Chen, M.-H. et al. Cilium-independent regulation of Gli protein function by Sufu in Hedgehog signaling is evolutionarily conserved. Genes Dev. 23, 1910–1928 (2009).

44. Haycraft, C. J. et al. Gli2 and Gli3 localize to cilia and require the intraflagellar transport protein polaris for processing and function. PLoS Genet. 1, e53 (2005).

45. Kim, J., Kato, M. & Beachy, P. A. Gli2 trafficking links Hedgehog-dependent activation of Smoothened in the primary cilium to transcriptional activation in the nucleus. Proc. Natl. Acad. Sci. U. S. A. 106, 21666–21671 (2009).

46. Dessauer, C. W. Adenylyl cyclase--A-kinase anchoring protein complexes: the next dimension in cAMP signaling. Mol. Pharmacol. 76, (2009).

47. Moore, B. S. et al. Cilia have high cAMP levels that are inhibited by Sonic Hedgehog-regulated calcium dynamics. Proc. Natl. Acad. Sci. U. S. A. 113, 13069–13074 (2016).

48. Somatilaka, B. N. et al. Ankmy2 Prevents Smoothened-Independent Hyperactivation of the Hedgehog Pathway via Cilia-Regulated Adenylyl Cyclase Signaling. Dev. Cell 54, 710–726.e8 (2020).

49. Jiang, J. Y., Falcone, J. L., Curci, S. & Hofer, A. M. Direct visualization of cAMP signaling in primary cilia reveals up-regulation of ciliary GPCR activity following Hedgehog activation. Proceedings of the National Academy of Sciences 116, 12066–12071 (2019).

50. Smith, F. D. et al. Local protein kinase A action proceeds through intact holoenzymes. Science 356, 1288–1293 (2017).

51. May, E. A. et al. Time-resolved proteomics profiling of the ciliary Hedgehog response. J. Cell Biol. 220, e202007207 (2021).

52. Calebiro, D. et al. Persistent cAMP-signals triggered by internalized G-protein-coupled receptors. PLoS Biol. 7, (2009).

53. Crilly, S. E. & Puthenveedu, M. A. Compartmentalized GPCR Signaling from Intracellular Membranes. J. Membr. Biol. 254, (2021).

54. Irannejad, R. et al. Conformational biosensors reveal GPCR signalling from endosomes. Nature 495, (2013).

55. Vilardaga, J.-P., Jean-Alphonse, F. G. & Gardella, T. J. Endosomal generation of cAMP in GPCR signaling. Nat. Chem. Biol. 10, 700–706 (2014).

56. Ogden, S. K. et al. G protein Gαi functions immediately downstream of Smoothened in Hedgehog signalling. Nature 456, 967–970 (2008).

57. Ayers, K. L. & Thérond, P. P. Evaluating Smoothened as a G-protein-coupled receptor for Hedgehog signalling. Trends Cell Biol. 20, 287–298 (2010).

58. Happ, J. T. et al. A PKA inhibitor motif within SMOOTHENED controls Hedgehog signal transduction. Nat. Struct. Mol. Biol. 29, 990–999 (2022).

59. Stubbs, T., Bingman, J. I., Besse, J. & Mykytyn, K. Ciliary signaling proteins are mislocalized in the brains of Bardet-Biedl syndrome 1-null mice. Frontiers in cell and developmental biology 10, (2023).

60. Badgandi, H. B., Hwang, S. H., Shimada, I. S., Loriot, E. & Mukhopadhyay, S. Tubby family proteins are adapters for ciliary trafficking of integral membrane proteins. J. Cell Biol. 216, (2017).

61. Sheu, S. H. et al. A serotonergic axon-cilium synapse drives nuclear signaling to alter chromatin accessibility. Cell 185, (2022).

62. Chou, C.-H. et al. Bisdemethoxycurcumin Promotes Apoptosis and Inhibits the Epithelial-Mesenchymal Transition through the Inhibition of the G-Protein-Coupled Receptor 161/Mammalian Target of Rapamycin Signaling Pathway in Triple Negative Breast Cancer Cells. J. Agric. Food Chem. 69, 14557–14567 (2021).

63. Bock, A. et al. Optical Mapping of cAMP Signaling at the Nanometer Scale. Cell 182, (2020).

64. Zhang, J. Z. et al. Phase Separation of a PKA Regulatory Subunit Controls cAMP Compartmentation and Oncogenic Signaling. Cell 182, (2020).

65. Ring, A. M. et al. Adrenaline-activated structure of β2-adrenoceptor stabilized by an engineered nanobody. Nature 502, 575–579 (2013).

66. Staus, D. P. et al. Sortase ligation enables homogeneous GPCR phosphorylation to reveal diversity in β-arrestin coupling. Proc. Natl. Acad. Sci. U. S. A. 115, 3834–3839 (2018).

67. Faust, B. et al. Autoantibody mimicry of hormone action at the thyrotropin receptor. Nature 609, 846–853 (2022).

68. Zheng, S. Q. et al. MotionCor2: anisotropic correction of beam-induced motion for improved cryo-electron microscopy. Nat. Methods 14, 331–332 (2017).

69. Punjani, A., Rubinstein, J. L., Fleet, D. J. & Brubaker, M. A. cryoSPARC: algorithms for rapid unsupervised cryo-EM structure determination. Nat. Methods 14, 290–296 (2017).

70. Asarnow, D., Palovcak, E. & Cheng, Y. asarnow/pyem: UCSF pyem v0.5. (2019). doi:10.5281/zenodo.3576630.

71. Pettersen, E. F. et al. UCSF ChimeraX: Structure visualization for researchers, educators, and developers. Protein Sci. 30, 70–82 (2021).

72. Zivanov, J. et al. New tools for automated high-resolution cryo-EM structure determination in RELION-3. Elife 7, (2018).

73. Grant, T., Rohou, A. & Grigorieff, N. cisTEM, user-friendly software for single-particle image processing. Elife 7, (2018).

74. Jumper, J. et al. Highly accurate protein structure prediction with AlphaFold. Nature 596, 583–589 (2021).

75. Adams, P. D. et al. PHENIX: a comprehensive Python-based system for macromolecular structure solution. Acta Crystallogr. D Biol. Crystallogr. 66, 213–221 (2010).

76. Emsley, P. & Cowtan, K. Coot: model-building tools for molecular graphics. Acta Crystallogr. D Biol. Crystallogr. 60, 2126–2132 (2004).

77. Schüttelkopf, A. W. & van Aalten, D. M. F. PRODRG: a tool for high-throughput crystallography of protein-ligand complexes. Acta Crystallogr. D Biol. Crystallogr. 60, 1355–1363 (2004).

78. Darbandi-Tonkabon, R. et al. Photoaffinity labeling with a neuroactive steroid analogue. 6-azi-pregnanolone labels voltage-dependent anion channel-1 in rat brain. J. Biol. Chem. 278, 13196–13206 (2003).

79. Zhang, L. & Hermans, J. Hydrophilicity of cavities in proteins. Proteins 24, 433–438 (1996).

80. Qu, C. et al. Ligand recognition, unconventional activation, and G protein coupling of the prostaglandin E_2_ receptor EP2 subtype. Sci Adv 7, (2021).

81. Lomize, M. A., Lomize, A. L., Pogozheva, I. D. & Mosberg, H. I. OPM: orientations of proteins in membranes database. Bioinformatics 22, 623–625 (2006).

82. Betz, R. Dabble. (2017). doi:10.5281/zenodo.836914.

83. Huang, J. et al. CHARMM36m: an improved force field for folded and intrinsically disordered proteins. Nat. Methods 14, 71–73 (2017).

84. Klauda, J. B. et al. Update of the CHARMM all-atom additive force field for lipids: validation on six lipid types. J. Phys. Chem. B 114, 7830–7843 (2010).

85. D.A. Case, H.M. Aktulga, K. Belfon, I.Y. Ben-Shalom, J.T. Berryman, S.R. Brozell, D.S. Cerutti, T.E. Cheatham, III, G.A. Cisneros, V.W.D. Cruzeiro, T.A. Darden, R.E. Duke, G. Giambasu, M.K. Gilson, H. Gohlke, A.W. Goetz, R. Harris, S. Izadi, S.A. Izmailov, K. Kasavajhala, M.C. Kaymak, E. King, A. Kovalenko, T. Kurtzman, T.S. Lee, S. LeGrand, P. Li, C. Lin, J. Liu, T. Luchko, R. Luo, M. Machado, V. Man, M. Manathunga, K.M. Merz, Y. Miao, O. Mikhailovskii, G. Monard, H. Nguyen, K.A. O’Hearn, A. Onufriev, F. Pan, S. Pantano, R. Qi, A. Rahnamoun, D.R. Roe, A. Roitberg, C. Sagui, S. Schott-Verdugo, A. Shajan, J. Shen, C.L. Simmerling, N.R. Skrynnikov, J. Smith, J. Swails, R.C. Walker, J. Wang, J. Wang, H. Wei, R.M. Wolf, X. Wu, Y. Xiong, Y. Xue, D.M. York, S. Zhao, and P.A. Kollman. Amber21. University of California, San Francisco (2022).

86. Salomon-Ferrer, R., Götz, A. W., Poole, D., Le Grand, S. & Walker, R. C. Routine Microsecond Molecular Dynamics Simulations with AMBER on GPUs. 2. Explicit Solvent Particle Mesh Ewald. J. Chem. Theory Comput. 9, 3878–3888 (2013).

87. Hopkins, C. W., Le Grand, S., Walker, R. C. & Roitberg, A. E. Long-Time-Step Molecular Dynamics through Hydrogen Mass Repartitioning. J. Chem. Theory Comput. 11, 1864–1874 (2015).

88. Roe, D. R. & Cheatham, T. E., 3rd. PTRAJ and CPPTRAJ: Software for Processing and Analysis of Molecular Dynamics Trajectory Data. J. Chem. Theory Comput. 9, 3084–3095 (2013).

89. Humphrey, W., Dalke, A. & Schulten, K. VMD: visual molecular dynamics. J. Mol. Graph. 14, 33–8, 27–8 (1996).

90. Hunter, J. D. Matplotlib: A 2D Graphics Environment. Comput. Sci. Eng. 9, 90–95 (2007).

91. Norman, R. X. et al. Tubby-like protein 3 (TULP3) regulates patterning in the mouse embryo through inhibition of Hedgehog signaling. Hum. Mol. Genet. 18, 1740–1754 (2009).

