## Supplementary Information for "GPR161 structure uncovers the redundant role of sterol-regulated ciliary cAMP signaling in the Hedgehog pathway"

**SUPPLEMENTARY INFORMATION for Hoppe et al:**

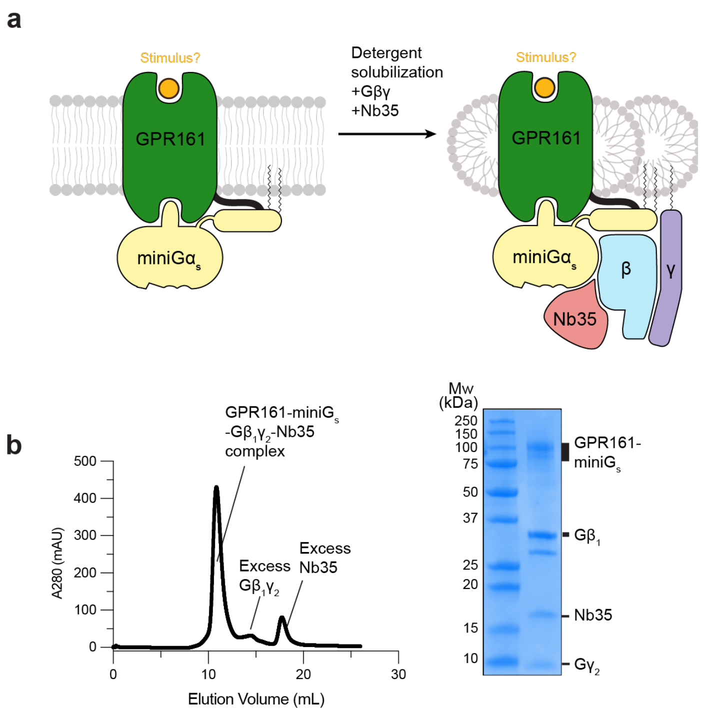


**Supplementary Figure 1: Biochemical preparation of GPR161-miniG$_s$ complex. a)** Cartoon depiction of GPR161 stabilization, solubilization, and purification. **b)** Size-exclusion chromatogram (left) and SDS-PAGE gel (right) of purified GPR161-G$_s$ complex with Nb35.

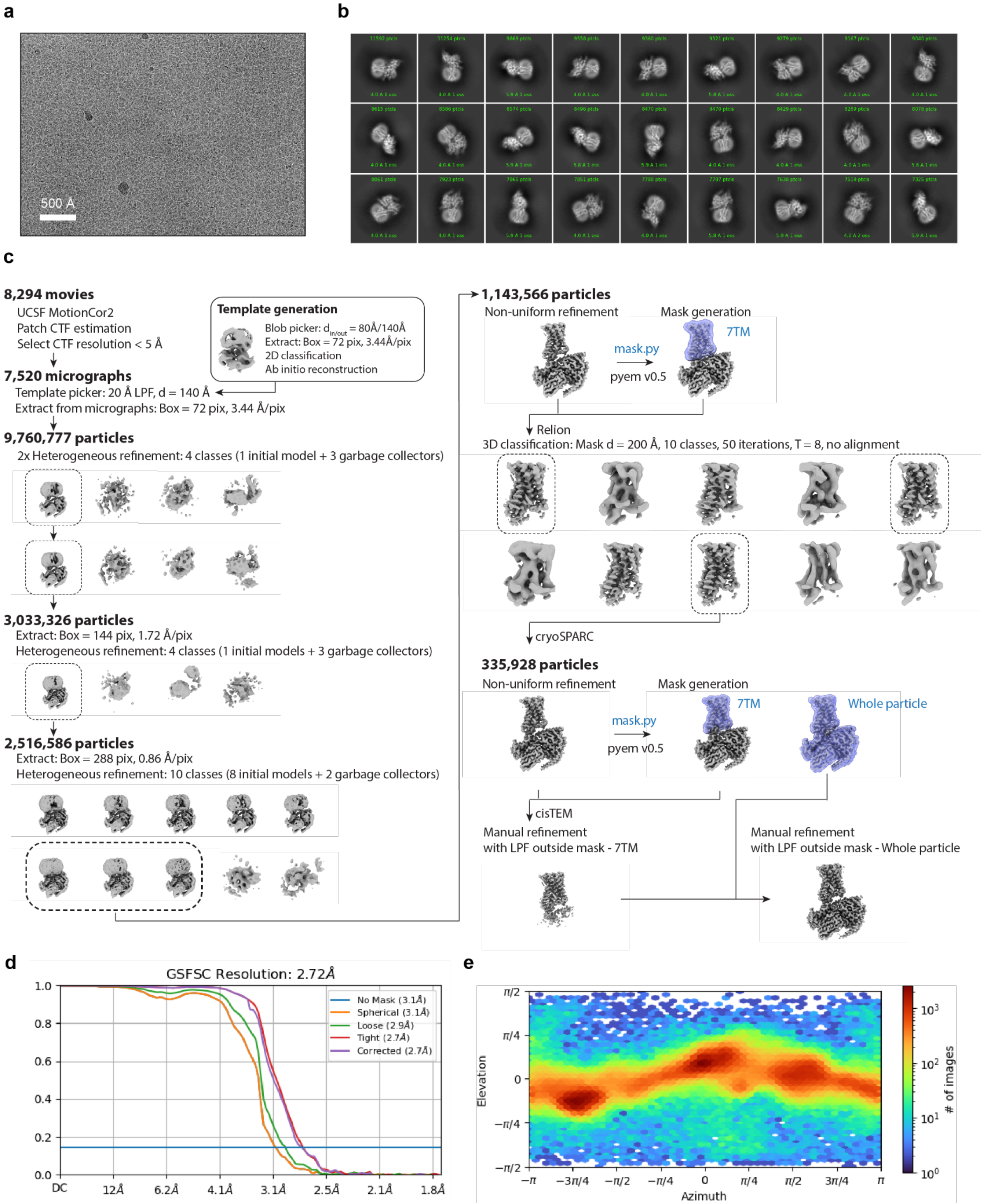


**Supplementary Figure 2: Cryogenic electron microscopy processing of GPR161. a)** A representative motion-corrected cryogenic electron microscopy (cryo-EM) micrograph obtained from a Titan Krios microscope (n = 8,294)**. b)** A subset of highly populated, reference-free 2D-class averages. **c)** Schematic showing the cryo-EM data processing workflow. Initial processing was performed using UCSF MotionCor2 and cryoSPARC. Particles were transferred using the pyem script package to RELION for alignment-free 3D classification. Finally, particles were processed in cisTEM using the manual refinement job type with a 7TM mask followed by a full particle mask. Dashed boxes indicated selected classes. **d)** Gold-standard Fourier Shell Correlation (GSFSC) curve for final refined and sharpened map computed in cryoSPARC. **e)** Euler angle distribution of final refined map computed in cryoSPARC.

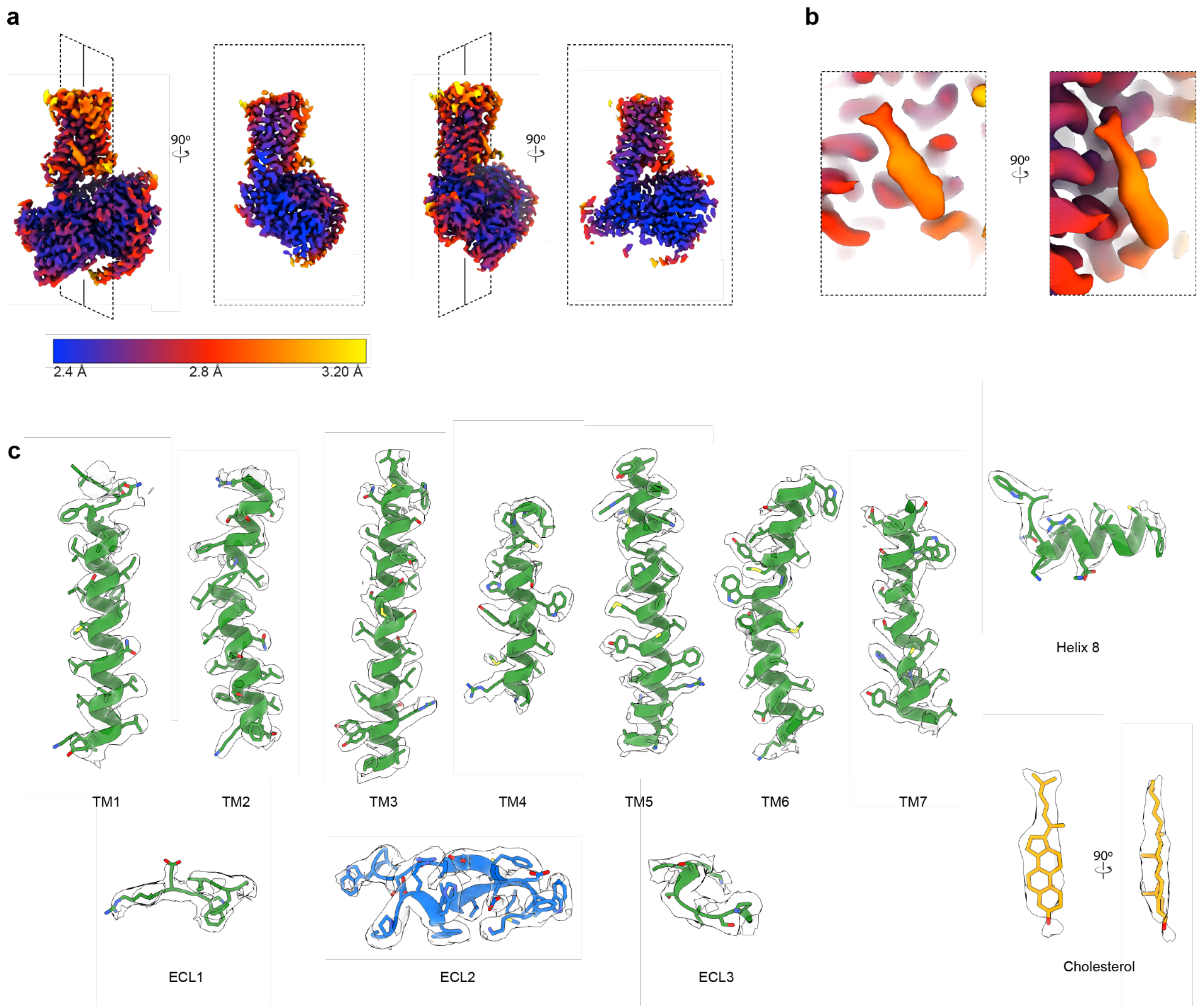


**Supplementary Figure 3: Cryo-EM local density. a)** Orthogonal views of local resolution for the sharpened, final map of GPR161-$G_s$ complex computed with local resolution in cryoSPARC. **b)** Close up of local resolution for sterol density. **c)** Isolated cryo-EM densities from the unsharpened, final map of GPR161 complex. Shown are the transmembrane (TM) helices, extracellular loops, and cholesterol-like density.

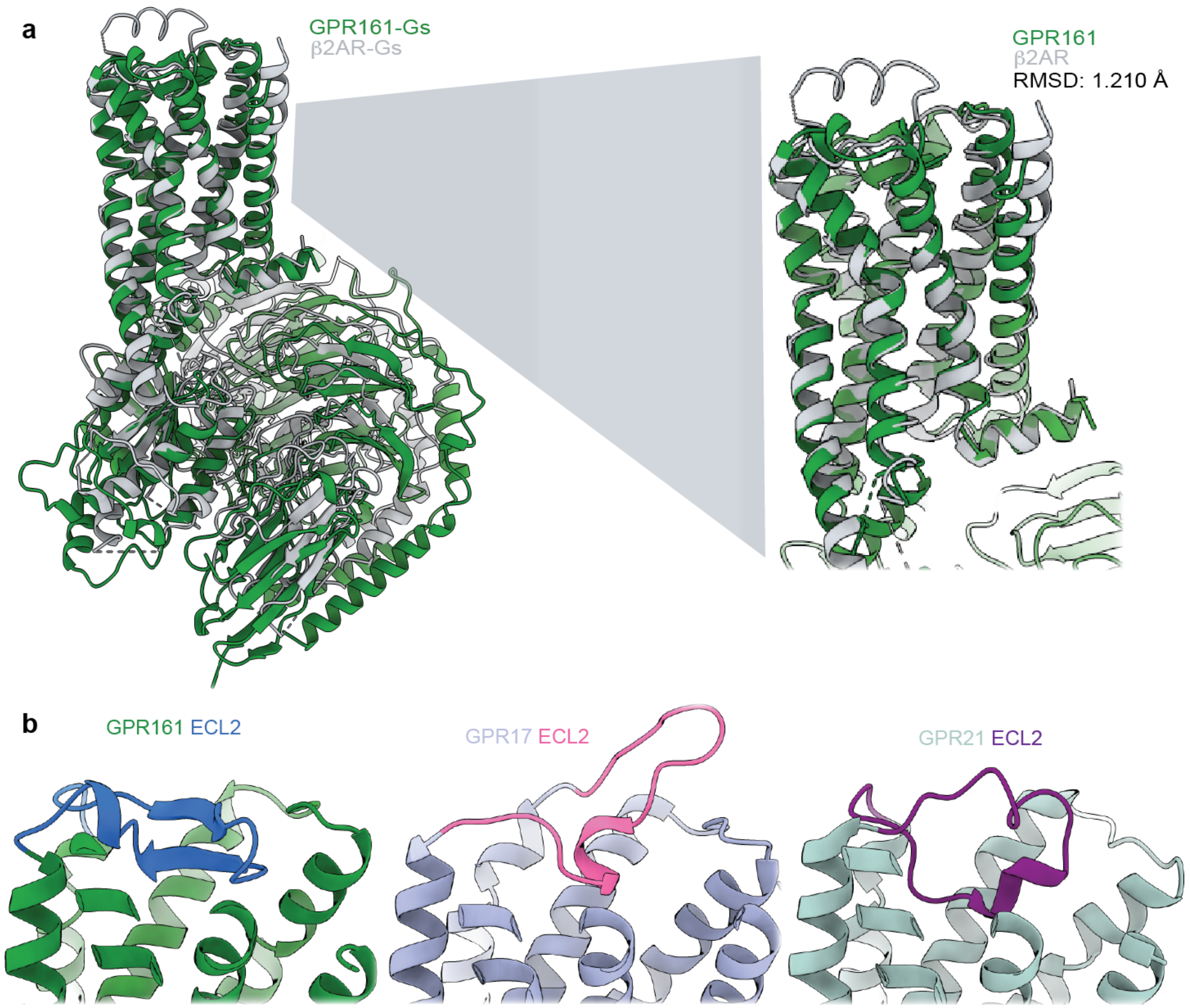


**Supplementary Figure 4: Comparison to additional GPCR structures. a)** Structural comparison of GPR161 heterotrimer complex and $\beta_2$AR heterotrimer complex (PDB ID: 3SN6 [65]). GPR161 has the same hallmarks of GPCR activation as the prototypical receptor, $\beta_2$AR **b)** Structural comparison of GPR161 to other orphan GPCRs with self-activating ECL2, including GPR17 (PDB ID: 7Y89) and GPR21 (PDB ID: 8HMV)[31,32]. The cis-interaction of ECL2 with the canonical ligand-binding site is seen across self-activating orphan GPCRs but the precise loop conformation changes between receptors.

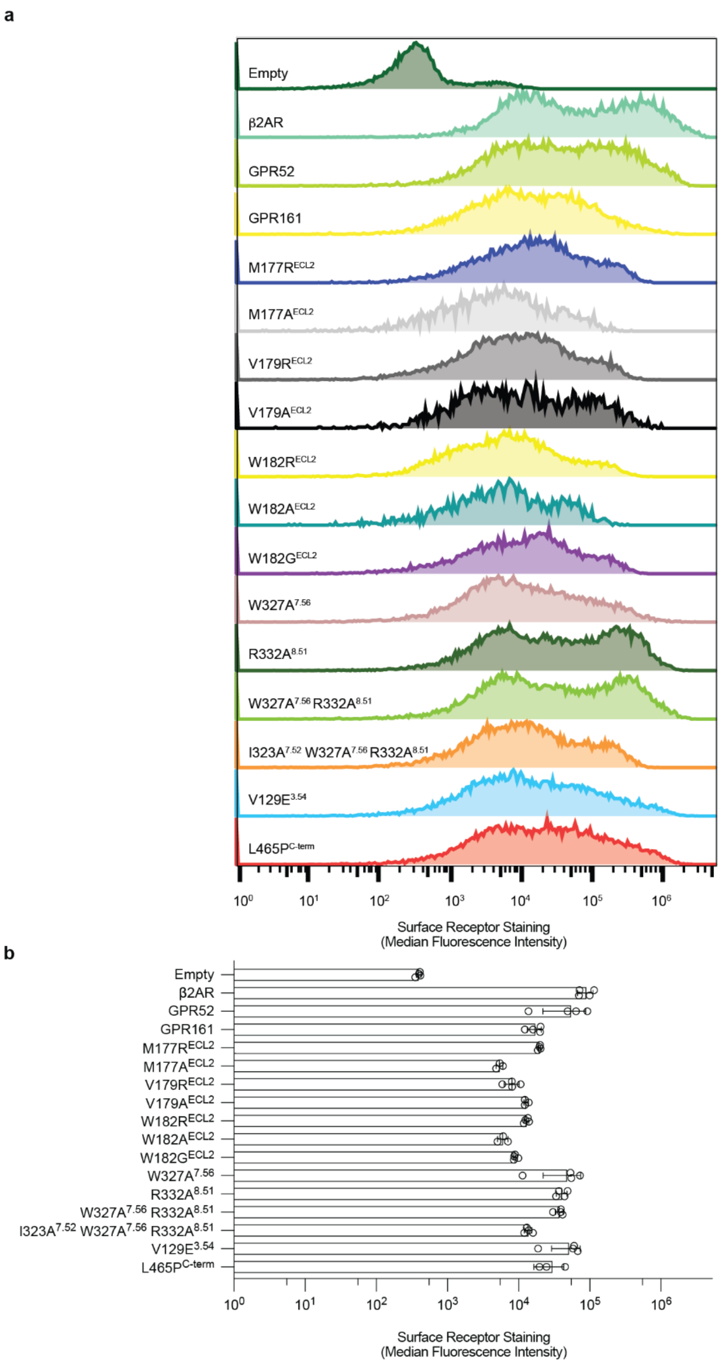


**Supplementary Figure 5: Surface expression of GPR161 mutants. a)** Representative flow cytometry surface expression histograms for receptors and mutants used in cell-based assays. **b)** Surface expression of receptors and mutants quantified by anti-FLAG-A647 median fluorescence intensity ± sd from n = 3-4 biologically independent samples.

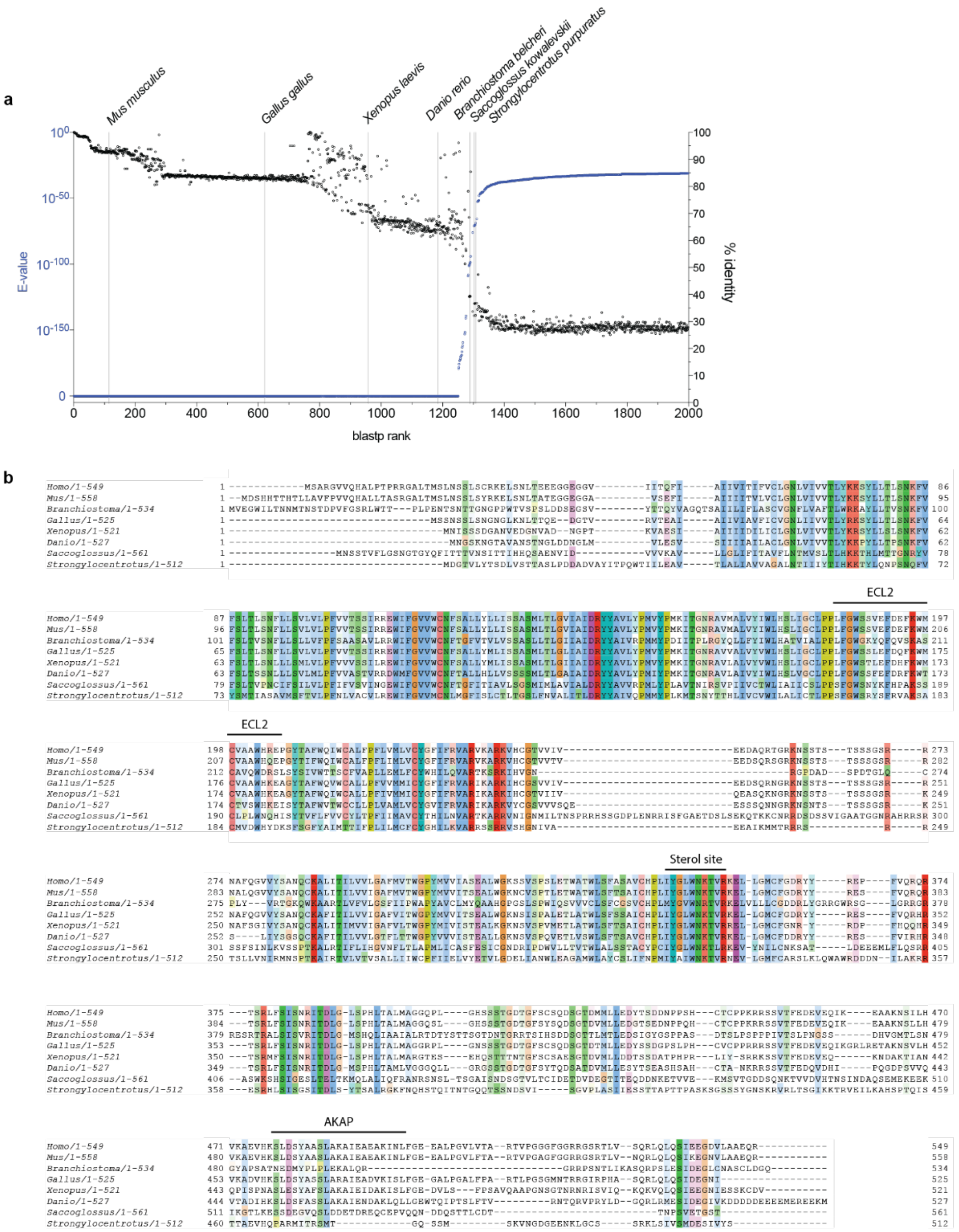


**Supplementary Figure 6: Phylogenetic analysis of GPR161. a)** BLAST search results for Human *GPR161* (Uniprot: Q8N6U8). Sequences are plotted from highest confidence (E-Value) and highest sequence identity (% identity) to lowest. Representative organisms spanning the full range of homologous GPR161 sequences are listed. **b)** Full sequence alignment of eight *GPR161* model organism sequences identified in BLAST search.

**a**

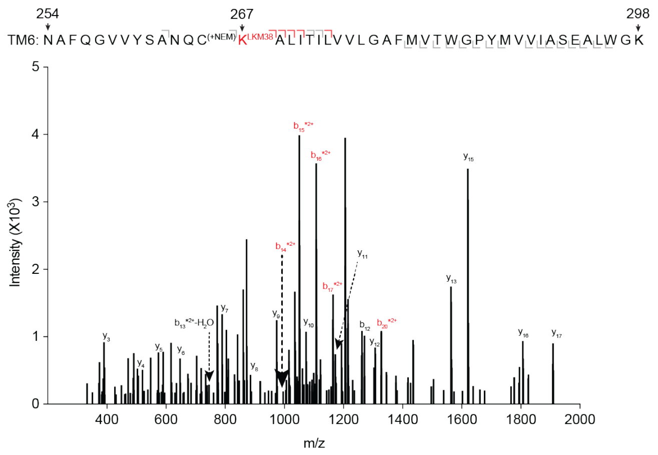


**b**

GPR161

>DYKDDDDASIDMSLNSSLSCRKELSNLTEEEGGEGGVIITQFIAIIVITIFVCLGNLVIV-
VTLYKKSYLLTLSNKFVFSLTLSNFLLSVLVLPFVVTSSIRREWIFGVVWCNFSALLYLLISSASMLTLGVIAIDRYYAVLYPMVYPMKITGNRAVMALVYIWLHSLIGCLPPLF
GWSSVEFDEFKWMCVAAWHREPGYTAFWQIWCALFPFLVMLVCYGFIFRVARVKARKVHCGTVVIVEEDAQRTGRKNSSTSTSSSGSRRNAFQGVVYSANQCKALIT
ILVVLGAFMVTWGPYMVVIASEALWGKSSVSPSLETWATWLSFASAVCHPLIYGLWNKTVRKELLGMCFGDRYYREPFVQRQRTSRLFSISNRITDLGLSPHLTALMAG
GQPLGHSSSTGDTGFSCSQDSGTDMMLLEDYTSDDNPPSHCTCPPKRRSSVTFEDEVEQIKEAAKNSILHVKAEVHKSLDSYAASLAKAIEAEAKINLFGEEALPGVLV
TARTVPGGGFGGRRGSRTLVSQRLQLQSIEEGDVLAAEQR

miniGs
>GGSLEVLFQGPSGGSGNSKTEDQRNEEKAQREANKKIEKQLQKDKQVYRATHRLLLL-
GADNSGKSTIVKQMRILHGGSGGSGGTSGIFETKFQVDKVNFHMFDVGGQRDERRKWIQCFNDVTAIIFVVDSSDYNRLQEALNLFKSIWNNRWLRTISVILFLNKQDL
LAEKVLAGKSKIEDYFPEFARYTTPEDATPEPGEDPRVTRAKYFIRDEFLRISTASGDGRHYCYPHFTCAVDTENARRIFNDCRDIIQRMHLRQYELL

Underlining: Transmembrane domains
Grey highlights: Glycosylation sequence
Red font: Peptides identified with tandem MS analysis

**Supplementary Figure 7: Photolabeling with LKM238 and mass spectrometry sequence coverage. a)** Product ion spectrum of LKM238-labeled GPR161-miniG$_s$ with peptides mapped to TM6. This peptide is modified with a mass consistent with LKM238 at position K267$^{6.32}$. Red brackets and peaks indicate product ions that contain the LKM238 adduct. **b)** Mass spectrometric sequence coverage of GPR161-miniG$_s$. Underlined segments indicate transmembrane spanning helices, red font indicates peptides identified by tandem MS analysis and gray font indicates glycosylation sites.

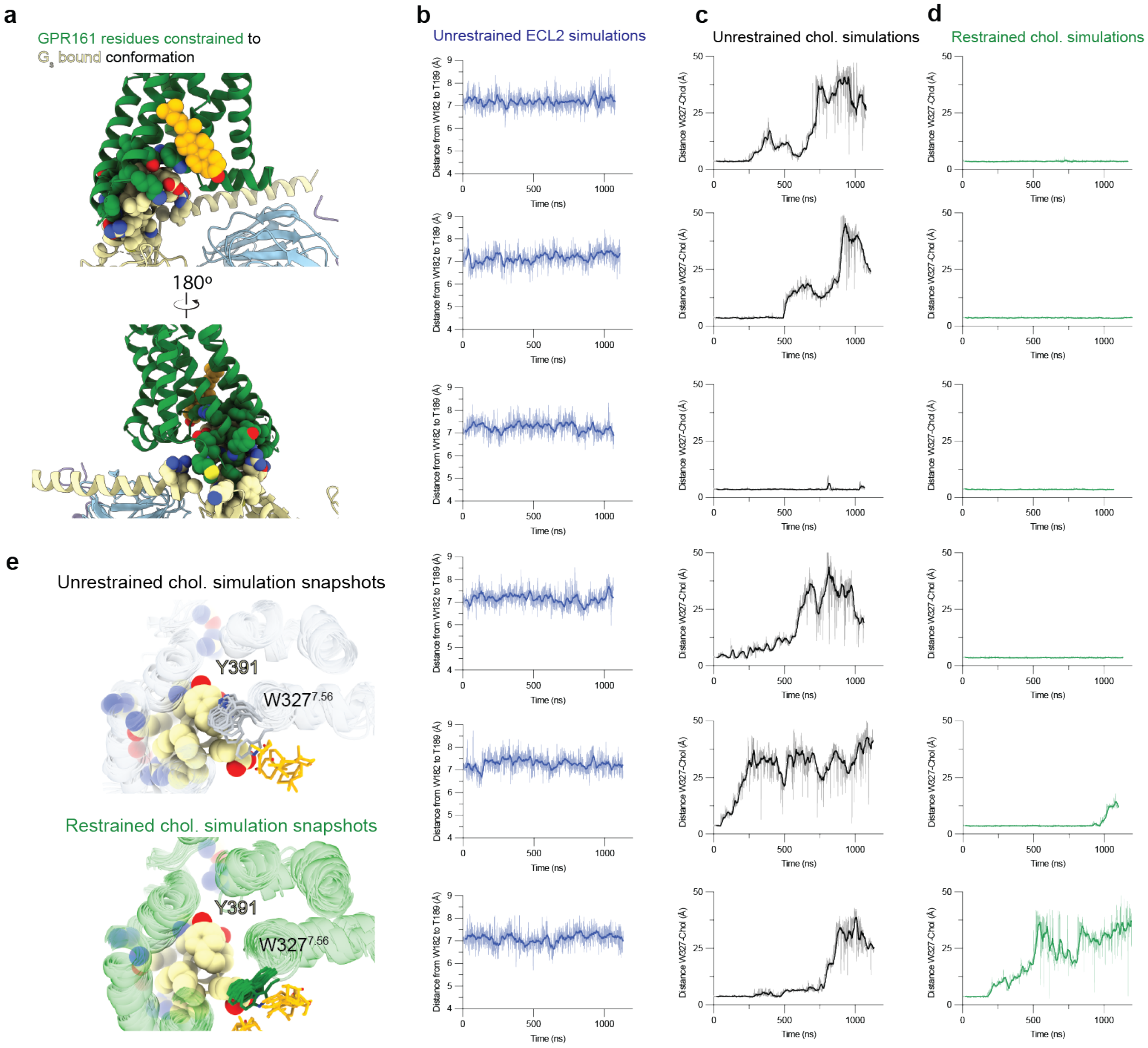


**Supplementary Figure 8: GPR161 molecular dynamics simulation trajectories. a)** Structural diagrams of the GPR161 residues that contact $G_s$, which are restrained to their $G_s$ bound conformation during restrained simulations. **b)** Independent trajectories for unrestrained simulations quantifying all atoms distance from $W182^{ECL2}$ to $T189^{5.39}$ . **c)** Independent trajectories for cholesterol-bound GPR161 without G protein-contacts restrained. All non-hydrogen atom distance from $W327^{7.56}$ to cholesterol is plotted. **d)** Independent trajectories for cholesterol-bound GPR161 with G protein-contacts restrained. All non-hydrogen atom distance from $W327^{7.56}$ to cholesterol is plotted. **e)** Simulation snapshots of $W327^{7.56}$ inward flip, which removes a key contact for cholesterol and occludes binding of the C-terminal α-helix of $G_s$.

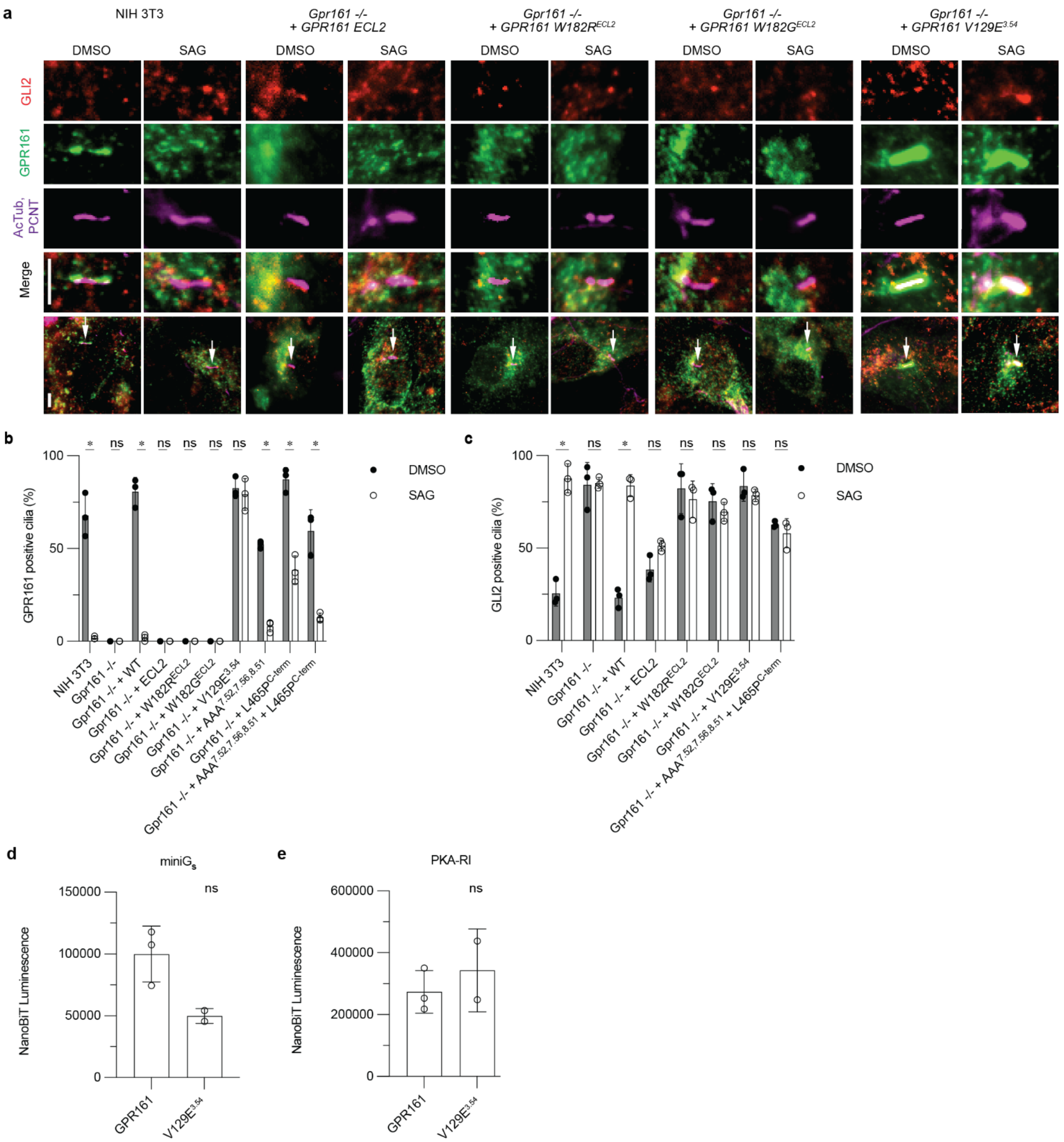


**Supplementary Figure 9: GPR161 localization and repression of ciliary trafficking of GLI2 for ECL2 mutants and V129E$^{3.54}$. a)** Representative images of the effect of site-directed mutagenesis of GPR161 on ciliary localization and GLI2 repression in ciliary tips in NIH 3T3 cells for ECL2 mutants and V129E$^{3.54}$. NIH 3T3 Flp-In CRISPR based *Gpr161$^{-/-}$* cells stably expressing untagged mouse wild-type or *GPR161* mutants were starved for 24 hr upon confluence and were treated for further 24 hr ± SAG (500 nM). After fixation, cells were immunostained with anti-GLI2 (red), anti-GPR161 (green), anti-acetylated, and centrosome (AcTub; PCNT purple) antibodies. Whole cell images with an arrow indicating imaged cilia. Scale bar, 5 μm. **b)** Quantification of GPR161 positive cilia indicating trafficking and egress of

GPR161 from cilia in the pathway off and on state, respectively. ECL2 mutants do not traffic to cilia suggesting impaired biogenesis. GPR161-V129E$^{3.54}$ does not egress from cilia following pathway activation and GPR161-L465P$^{C\text{-term}}$ has reduced egress compared to GPR161. All other mutants traffic similar to GPR161. Quantification of GPR161 positive cilia are shown from 3 independent experiments from images taken from 2-3 different regions/experiment and counting 15-30 cells/region. Data are mean ± s.d. (**P* < 0.05; ns, not significant; two-way ANOVA followed by Šidák's multiple comparison tests). **c)** Quantification GLI2 positive cilia indicating Hedgehog pathway activation. ECL2 mutants and GPR161-V129E$^{3.52}$ do not rescue, similar to *Gpr161*$^{-/-}$. Quantification of GLI2 positive cilia are shown from 3 independent experiments from images taken from 2-3 different regions/experiment and counting 15-30 cells/region. Data are mean ± s.d. (**P* < 0.05; ns, not significant; two-way ANOVA followed by Šidák's multiple comparison tests). **d)** GPR161-V129E$^{3.54}$ has reduced recruitment of miniG$_s$ compared to WT. Data are mean ± s.d., n=2-3 biologically independent samples (**P* < 0.05; ns, not significant; one-way ANOVA followed by Dunnett's multiple comparison tests). **e)** GPR161-V129E$^{3.54}$ has similar recruitment of PKA-RI compared to GPR161. Nanoluc complementation assay for receptor recruitment of miniG$_s$. Both GPR161 and GPR161-L465P$^{C\text{-term}}$ constitutively recruit miniG$_s$ while GPR161-AAA$^{7.52,\ 7.56,\ 8.51}$ does not. Data are mean ± s.d., n=2-3 biologically independent samples (**P* < 0.05; ns, not significant; one-way ANOVA followed by Dunnett's multiple comparison tests)..

**Supplementary Table 1: Cryo-EM data collection and model statistics.**

| | **GPR161-$G_s$** |
|---|---|
| EMDB: Full map | EMD-40603 |
| RCSB PDB: Model | 8SMV |
| **Data collection** | |
| Microscope | Thermo Scientific Krios G3i |
| Detector | Gatan K3 with Gatan BioQuantum Energy filter |
| Voltage (kV) | 300 |
| Magnification | 105,000 |
| Defocus range (µm) | -0.8 to -2.2 |
| Pixel size, physical (Å) | 0.86 |
| Total exposure ($e^-/Å^2$) | 50.7 |
| Frame exposure ($e^-/Å^2$/frame) | 0.845 |
| Images, number of | 8,294 |
| Frames/image, number of | 60 |
| Initial particles, number of | 9,760,777 |
| Final particles, number of | 335,928 |
| Symmetry imposed | C1 |
| Map sharpening, *B* factor ($Å^2$) | |
| Full map | -90 |
| Map resolution, masked (Å) | |
| Full map | 2.72 |
| FSC threshold | 0.143 |
| **Refinement** | |
| Initial model used (AlphaFold code) | Q8N6U8 |
| Model resolution (Å) | 2.72 |
| Model composition | |
| Chains | 6 |
| Non-hydrogen atoms | 8,169 |
| Protein residues | 1,034 |
| Ligands | 1 |
| *B* factors ($Å^2$) | |
| Receptor | 45.0 |
| Ligand | 53.52 |
| G protein with Nb35 | 21.58 |
| R.m.s. deviations | |
| Bond length (Å) | 0.004 |
| Bond angles (°) | 1.033 |
| Validation | |
| MolProbity score | 1.12 |
| Clash score | 1.85 |
| EMRinger score | 3.47 |
| Rotamer outliers (%) | 0.00 |
| Ramachandran plot | |
| Favored (%) | 96.96 |
| Allowed (%) | 3.04 |
| Disallowed (%) | 0.00 |
